# Early life stress influences epilepsy outcomes in mice

**DOI:** 10.1101/2024.09.09.612052

**Authors:** Emanuel M Coleman, Maya White, Pantelis Antonoudiou, Grant L Weiss, Garrett Scarpa, Bradly Stone, Jamie Maguire

**Author notes:** Corresponding Author: Jamie Maguire Tufts University School of Medicine Neuroscience Department 136 Harrison Ave. Boston, MA 02111.

## Abstract

Stress is a common seizure trigger that has been implicated in worsening epilepsy outcomes. The neuroendocrine response to stress is mediated by the hypothalamic-pituitary-adrenal (HPA) axis and HPA axis dysfunction worsens epilepsy outcomes, increasing seizure burden, behavioral comorbidities, and risk for sudden unexpected death in epilepsy (SUDEP) in mice. Early life stress (ELS) reprograms the HPA axis into adulthood, impacting both the basal and stress-induced activity. Thus, we propose that ELS may influence epilepsy outcomes by influencing the function of the HPA axis. To test this hypothesis, we utilized the maternal separation paradigm and examined the impact on seizure susceptibility. We show that ELS exerts a sex dependent effect on seizure susceptibility in response to acute administration of the chemoconvulsant, kainic acid, which is associated with an altered relationship between seizure activity and HPA axis function. To further examine the impact of ELS on epilepsy outcomes, we utilized the intrahippocampal kainic acid model of chronic epilepsy in mice previously exposed to maternal separation. We find that the relationship between corticosterone levels and the extent of epileptiform activity is altered in mice subjected to ELS. We demonstrate that ELS impacts behavioral outcomes associated with chronic epilepsy in a sex-dependent manner, with females being more affected. We also observe reduced mortality (presumed SUDEP) in female mice subjected to ELS, consistent with previous findings suggesting a role for HPA axis dysfunction in SUDEP risk. These data demonstrate for the first time that ELS influences epilepsy outcomes and suggest that previous life experiences may impact the trajectory of epilepsy.

## Introduction

Stress is a common seizure trigger, and cortisol levels are basally elevated in people with epilepsy (PWE) which increase postictally, correlating with seizure severity (for review see ^1^).

The impact of stress on epilepsy outcomes is thought to be mediated by the actions of neuroendocrine stress mediators, such as cortisol and corticotropin releasing hormone (CRH), both of which have been shown to exert proconvulsant effects, exacerbate neuropathology, increase comorbid behavioral deficits, and accelerate epileptogenesis and disease progression (^2–5^ (for review see ^6,7^).

The physiological response to stress is mediated by the hypothalamic-pituitary-adrenal (HPA) axis. In response to stress or seizures, CRH is released from the paraventricular nucleus of the hypothalamus (PVN), which governs HPA axis activity, and sequentially triggers the release of adrenocorticotropin hormone (ACTH) from the anterior pituitary gland, and cortisol from the adrenal glands (corticosterone in mice). We previously demonstrated that HPA axis dysfunction worsens epilepsy outcomes, increasing seizure susceptibility/frequency, vulnerability to comorbid behavioral deficits, and the incidence of sudden unexpected death in epilepsy (SUDEP) in chronically epileptic mice ^2,3^.

Early life stress (ELS) has been demonstrated to reprogram HPA axis function (for review see ^8^. The perinatal period encompasses a critical developmental window involving extensive plasticity which is sensitive to perturbations by external forces, such as stress (for review see ^9,10^). Abundant evidence demonstrates that early life stress during this sensitive period can induce lasting effects on behavioral outcomes, impacting the ability to cope with stress and increasing the risk for psychiatric illnesses such as major depressive disorder (MDD) and posttraumatic stress disorder (PTSD). For example, adverse childhood events (ACEs) have been demonstrated to increase arousal ^11^, enhance negative valence processing ^12,13^, and induce cognitive deficits ^14,15^. Similar effects have been demonstrated in animal models, in which early life stress has been demonstrated to lead to deficits in behaviors, including increased avoidance behaviors, that persist into adulthood (for review see ^16–18^). These negative consequences of early life stress are thought to involve the long-term alterations in HPA axis function following ELS. Although numerous studies report that ELS induces HPA axis hyperexcitability, pathological HPA axis hypofunction has also been demonstrated ^19–21^ (for review see ^8^). Additionally, it has been suggested that ELS prepares an animal to respond to similar adversity later in life and produce a predictive adaptive response ^22^ (for review see ^23^).

Previous studies have demonstrated that early life stress impacts epileptogenesis (for review see ^24–26^. The majority of these studies, using a variety of early life stress paradigms, report an increased vulnerability to epilepsy following early life stress (for review see ^24,26^). It is important to note that previous studies in several models have demonstrated sex differences in epilepsy outcomes following early life stress. Many studies across models have demonstrated that males are more affected by early life stress with increased vulnerability to epilepsy than females ^27–30^; whereas, it has been shown the females are more impacted in the maternal separation model ^31,32^. However, the majority of studies have focused solely on the impact of early life stress on seizure outcomes and the impact on associated comorbidities remains less clear. Although one impactful study demonstrated that early life stress accelerated epileptogenesis and increased depression-like behaviors in chronically epileptic rats ^33^. Here we build on these foundational studies to further interrogate the impact of early life stress on epilepsy outcomes, focusing on psychiatric comorbidities and SUDEP risk. Interestingly, these previous studies have implicated HPA axis dysfunction in mediating the impact of early life stress on seizure outcomes (for review see ^23–26^). In fact, the sex differences in the impact of early life stress on the development of epilepsy was linked to a rise in corticosterone regardless of sex which correlated with the development of epilepsy in this model ^30,31,34^. Further, the impact of early life stress on seizure susceptibility and epileptogenesis is thought to be mediated by stress hormones and HPA axis dysfunction ^25,31^.

Given the demonstrated role of HPA axis dysfunction in worsening epilepsy outcomes ^2,35–37^ (for review see ^38^), we proposed that ELS may impact epilepsy outcomes through modulation of HPA axis activity. To test this hypothesis, we utilized a maternal separation paradigm to model early life stress and evaluated the impact on HPA axis function in relation to both acute and chronic seizure susceptibility, the impact on comorbid behavioral deficits in chronically epileptic mice, and SUDEP risk. Here, we demonstrate that ELS alters seizure susceptibility in a sex dependent manner in response to acute administration of the chemoconvulsant, kainic acid. Interestingly, we observe that ELS impacts the relationship between seizure-induced elevation in corticosterone levels compared to controls. Further, while there is no significant impact of ELS on seizure frequency in chronically epileptic mice, there is a correlation between corticosterone levels and seizure activity. Although we did not observe any differences in chronic seizure burden, our data demonstrates sex dependent impacts of ELS on behavioral deficits associated with epilepsy and mortality rate, consistent with previous work implicating HPA axis dysfunction in SUDEP risk. These data demonstrate for the first time that previous life experiences can impact the trajectory of epilepsy which may involve altered HPA axis function.

## Material and methods

### Animals

C57BL/6J mice were housed at Tufts University School of Medicine in a temperature and humidity-controlled environment, maintained on a 12:12 light/dark cycle (lights on at 7 a.m.) with *ad libitum* access to food and water. All animal procedures were handled according to the protocols approved by the Tufts University Institutional Animal Care and Use Committee (IACUC).

### Early life stress (ELS) paradigm

Mice were subjected to the maternal separation paradigm of early life stress as previously described ^39,40^. Timed pregnant C57BL/6J dams were purchased directly from Jackson Laboratory (Bar Harbor, ME) and delivered to Tufts University School of Medicine on day 16 of gestation (E16). Dams were singly housed and checked daily for litters, at which time they were randomly assigned to standard-rearing (Ctrl) or maternal separation (ELS) groups. ELS mice were subjected to 3hrs of daily maternal separation (9a-12p) for 15 days over postnatal day 1-21 (PND1-21). During separation, ELS dams were removed from their home cage and placed in a clean holding cage out of sight from the litter. Offspring were weaned on PND28 and left undisturbed until adulthood (>PND70). Ctrl mice were similarly handled without removal of the dam from the home cage.

### Stereotaxic surgery

Adult mice previously subjected to ELS or Ctrl rearing conditions were anesthetized with a ketamine/xylazine cocktail (90-120 mg/kg ketamine, 5-10 mg/kg xylazine; *i.p.*) and treated with sustained release buprenorphine (0.5-1 mg/kg; *s.c.*) prior to surgical procedures. For all electroencephalogram recordings (*acute-EEG* and *chronic-EEG*), mice were implanted with a prefabricated EEG headmount (Pinnacle Technology, cat. #8201) to measure electrographic epileptiform activity and record spontaneous seizures. The headmount was fixed to the skull with dental cement and four screws: one EEG lead was placed in each of the frontal and parietal cortex, and two screws were placed in the opposite hemisphere which serve as the reference and animal ground. Electroencephalogram recordings were collected at 4 KHz using a 100X gain preamplifier high pass filtered at 1 KHz (Pinnacle Technology, cat. #8292-SE) and tethered turnkey system (Pinnacle Technology, cat. #8200).

### Acute seizure susceptibility testing

Acute seizure susceptibility testing (*acute-EEG*) began 5-7 days post-surgery. Testing was performed in awake, behaving animals in response to acute treatment with the chemoconvulsant kainic acid (20 mg/kg KA, *i.p.*). Mice were habituated to the EEG recording chamber for 45 mins prior to KA administration and EEG recordings were acquired for 2 hrs following kainic acid administration. Percent electrographic epileptiform activity, the latency to the first seizure event and total time exhibiting abnormal electrographic epileptiform activity were measured over the 2 hr recording period. Abnormal electrographic activity was defined as paroxysmal activity having a sudden onset and an amplitude of at least 2.5x the standard deviation of the baseline and a consistent change in the power of the fast fourier transform of the EEG, including a change in the power and the frequency of activity over the course of the event. Abnormal electrographic activity, including ictal activity and rhythmic spiking lasting longer than 30 s were collectively defined as “epileptiform activity”. Latency to the first ictal event was defined as the time elapsed from the injection of KA to the start of the first identified electrographic seizure. For each mouse, percent epileptiform activity was calculated as:

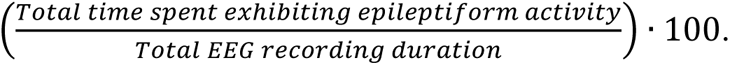

*Status epilepticus* (SE) was defined as unremitting epileptiform activity lasting longer than 5 minutes according to the clinical definition ^41,42^. The latency to the onset of SE was measured and the total time spent in SE was measured.

### Ventral intrahippocampal kainic acid (vIHKA) model of chronic epilepsy

To assess the impact of ELS in chronically epileptic mice, we employed the ventral intrahippocampal kainic acid (vIHKA) model. This protocol has been used by our laboratory and others to generate chronically epileptic mice ^2,43^. Briefly, mice were stereotaxically injected with kainic acid (100nL, 20mM) unilaterally into the ventral hippocampus (from Bregma in mm: A/P -3.6, M/L -2.8, D/V -2.8) using a 33-gauge Hamilton syringe at an infusion rate of 50nL/min. Following vIHKA infusion, a subset of mice were implanted with EEG headmounts to record spontaneous seizure activity. Continuous EEG recordings in chronically epileptic mice (*chronic-EEG*) began between 7-10 days post-surgery and continued uninterrupted for two weeks.

Seizures from EEG traces were automatically detected using a python application previously developed and validated in our laboratory (https://github.com/neurosimata/seizy) ^2,44^. Briefly, a thresholding method was applied to downsample traces to 100Hz and divide them into 5 second segments. Seizure segments were detected based on power (2-40 Hz) and line length of the downsampled EEG signal. Seizures were only included if two consecutive 5 s segments were classified as seizures (minimum 10 s length). Automatically detected seizures were manually verified and false positive seizures were discarded. Seizure frequency was calculated by dividing the total number of seizures a mouse had over the recording period by the total number of recording hours. The average seizure duration was calculated by taking the mean of seizure event durations for each mouse in each recording period. Seizure burden was then calculated as the seizure frequency times the seizure duration for each animal. The number of seizures detected by the Seizy application strongly correlated to manual scored data from 6 pseudo-randomly selected 24hr recordings (*R*^2^=0.95, *r*= 0.97, *p*=0.0011).

### Survival criteria

Survivability of all vIHKA Ctrl and ELS mice was monitored from the induction of SE until the start of behavioral testing. Any animals that died within the first three days post-surgery were excluded from the survival curve and resulting analyses. No deaths occurred during behavioral testing. All post-mortem animals that were found with characteristic hindlimb extention i.e. complete tonic extension were presumed to have died by SUDEP. Bodyweight was measured once at surgery and again at euthanasia to assess any changes in mass.

### Behavioral paradigms

Behavioral testing was conducted in chronically epileptic Ctrl and ELS mice beginning at 60 days following vIHKA infusion. Avoidance behaviors were tested using the open field test (OFT), light/dark box (L/D), and elevated plus maze (EPM) as previously described ^2,45,46^. The testing order was counterbalanced to avoid any order effects. Stress-induced helplessness was measured using the tail suspension test (TST) and anhedonia was evaluated using the sucrose preference test (SPT). The animal’s innate startle reflex to an unavoidable acoustic stimulus was tested using the acoustic startle response (ASR). Home cage locomotor activity (LA) was recorded over 72hrs.

#### Open field test (OFT)

Avoidance behaviors in the open field test were measured as previously described ^2,45,46^. Briefly, mice were placed in an open arena surrounded by a 40 cm x 40 cm photobeam frame with 16 x 16 equally spaced photocells (Hamilton-Kinder). Mice were placed individually into the center of the arena and beam breaks and movement were automatically detected using the Motor Monitor software (Hamilton-Kinder) over the 10 min testing period. The time spent and distance traveled in the center of the open field was quantified, as well as the total distance traveled during the testing period.

#### Light/dark box (LD Box)

Avoidance behaviors in the light/dark box were measured as previously described ^2,45,46^. Mice were placed in the dark chamber of a two-chamber light/dark box apparatus surrounded by a 22 cm x 43 cm photobeam frame with 8 equally spaced photocells (Hamilton-Kinder) and allowed to explore the two-chamber apparatus freely for a 10 min testing period. Beam breaks and movement were detected using the automated Motor Monitor software (Hamilton-Kinder). The time spent and distance traveled in both the light and dark chambers were measured, as well as the total distance traveled during the testing period.

#### Elevated plus maze (EPM)

Avoidance behaviors in the elevated plus maze were measured as previously described ^2,45,46^. Briefly, mice were placed in the intersection of a 78 cm x 78 cm x 78 cm elevated maze apparatus with 48 infrared photobeams (Hamilton-Kinder). The maze had an intersection zone (5 cm x 5 cm) separating all four arms (5 cm x 38 cm), two of which were enclosed by 15 cm walls. Beam breaks and movement were automatically detected through the Motor Monitor software (Hamilton-Kinder) over the 10 min testing period. The time spent and distance traveled in both the open and closed arms were measured.

#### Home cage locomotor activity testing (LA)

To test for relative hyperactivity and locomotor differences in chronically epileptic ELS and Ctrl mice, baseline locomotor activity was monitored in cages identical to their home cage surrounded by a 22 cm x 43 cm photobeam frame with 8 equally spaced photocells (Hamilton-Kinder). Beam breaks and movement were automatically detected using the Motor Monitor software (Hamilton-Kinder). Testing began at the same time for all mice (ZT 5) and lasted 72 hrs. The total distance traveled was measured for each mouse and analyzed in 6 hr bins to capture circadian locomotor patterns.

#### Sucrose preference test (SPT)

Similar to previous studies, anhedonia was measured through the sucrose preference test^2^. Mice were habituated to specialty cages that contained two identical bottles filled with water for 2 days prior to the testing period. On the first day of testing, mice were singly housed and given *ad libitum* access to two new bottles, one filled with water and the other filled with 2% sucrose (w/v). Bottles were weighed and their positions swapped daily to avoid place preference. The testing period lasted 5 days and each mouse’s sucrose preference score was calculated by dividing the daily change in the sucrose water bottle weight by the sum of the changes in the bottle weights for both the regular water and sucrose water (Δ sucrose/Δ sucrose + Δ reg. water).

#### Tail suspension test (TST)

Stress-induced helplessness was measured using the TST as previously described by our laboratory ^47^. Briefly, mice were suspended by their tails from a platform roughly 36 cm above the ground for 6 mins. Trials were video recorded and automatically scored using an unbiased in-house Mobility App (https://github.com/researchgrant/mobility-mapper.git) to measure the latency to immobility and the total time spent immobile. Limb movement (nose, paws, and tail) were calculated from positional data automatically detected using DeepLabCut. A neural network was then trained using a Long Short-Term Memory recurrent neural network architecture in the TensorFlow toolbox for Python to predict the mobility state of the animals based on videos scored by two different investigators. The mobility network’s time spent immobile correlated to manual scored data (*R*^2^=0.52, *r*= 0.71, *p*<0.0001).

#### Acoustic startle response (ASR)

The startle reflex in chronically epileptic ELS and Ctrl mice was measured in response to unavoidable acoustic stimuli. Briefly, startle testing was conducted in an SM100 startle reflex station (Hamilton-Kinder) housed within a sound-attenuated cabinet and equipped with built-in speakers. Mice were placed in the restraining chamber on a piezoelectric plate for the test while stimuli were generated and recorded with the Startle Monitor software (Hamilton-Kinder). The startle paradigm consisted of a 3 min acclimatization period followed by six pseudorandomized white noise bursts of 50ms at different intensities (95 dB / 100 dB / 110 dB, 2x each) at a randomized intertrial interval (ITI = 45-90 s, average = 70 s) to avoid habituation. The background level and recording window were set to 65 dB and 250 ms, respectively. The max, average, and first startle response (N) was measured for each mouse and averaged across intensities.

### Corticosterone ELISAs

Mice were anesthetized with isoflurane and euthanized by rapid decapitation for brain extraction and trunk blood collection. For *acute-EEG* mice, blood was collected 2 hrs after KA injections (20 mg/kg, *i.p.*). For chronically epileptic vIHKA mice, blood was collected at the end of behavioral testing after 24 hrs of habituation to the same room where euthanasia was performed. Blood collection occurred at the same time for all mice (ZT 3-4). Blood samples were placed on ice immediately after collection, allowed to reach room temperature, and centrifuged at 14K rpm for 15 mins at 4°C to isolate serum. Serum was stored at -80°C until use. To quantify corticosterone (CORT) levels from our serum samples, we used a corticosterone enzyme linked immunosorbent assay (ELISA) kit (Enzo Life Sciences, cat. #ADI-900-097). Each sample was run in duplicate and experimental samples were compared to a standard curve of known corticosterone concentrations.

### Immunohistochemistry

Brains were rapidly extracted, fixed using immersion fixation in 4% paraformaldehyde at 4°C overnight, and subsequently cryoprotected in 10-30% sucrose, embedded in Tissue-Tek O.C.T., and stored at -80°C until cryostat sectioning. Brains were coronally sectioned on a crysostat at 40 µm. For c-Fos staining, sections were blocked for 2 hrs in 5% normal goat serum at room temperature and incubated in primary anti-cFos (1:1000; Synaptic Systems, cat. # 226 008) for 2 days at 4°C. Sections were then incubated in a goat anti-rabbit Alexa Fluor 488 secondary (1:200; Invitrogen, cat. # A-11008) for 2 hrs at room temperature. Sections were mounted with a medium containing DAPI (Vector Laboratories) to enable identification of the region of interest, the paraventricular nucleus of the hypothalamus (PVN), based on anatomical structures. Stained sections were visualized using a Keyence BZ-X700 microscope and imaged at 10x for quantification of the number of c-Fos-positive neurons in the PVN. Representative slices of the PVN were imaged and analyzed for each animal.

Individual cells expressing c-Fos were detected and quantified using the included Cell Profiler pipelines. Briefly, the Minimum Cross-Entropy detection method with global thresholding was used to detect cells from the DAPI channel. The threshold correction factor was adjusted between 1.0 and 1.05 to manually adjust for variance in tissue thickness. A PVN mask was then used to select only neuronal cells within the region of interest (ROI). The percent c-Fos positive cell count in the manually-outlined ROI was then calculated as:

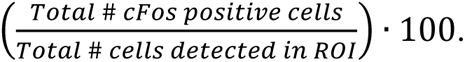

### Statistical analyses

Data was analyzed using GraphPad Prism 9. The details of all statistical tests can be seen in the statistical tables associated with each figure. When comparing two experimental groups, an unpaired *t*-test, Welch’s *t*-test or Mann-Whitney test was used to determine statistical significance. Multiple conditions were either compared using a two-way ANOVA with Tukey’s post-hoc multiple comparisons, or a mixed-effects analysis with Šidak’s post-hoc multiple comparisons. For *chronic-EEG* animals, correlational analyses were performed between seizure metrics and basal CORT levels. All data were tested for normalcy and outlier removal via the ROUT method (Q=5%). All *p*-values <0.05 were considered significant (# = *p* < 0.1; * = *p* < 0.05; ** = *p* < 0.01; *** = *p* < 0.001; **** = *p* < 0.0001).

### Multiple linear regression

To model the relationship between seizure and behavioral outcomes in the chronic vIHKA model of TLE, we performed a least-squares multiple linear regression on 22 *chronic-EEG* mice (6 Ctrl F, 5 ELS F, 6 Ctrl M, 5 ELS M) using GraphPad Prism 9. First, all features were z-normalized. Following normalization, highly correlated variables with r > 0.6 were removed, resulting in 11 features that were fed into the model. The equation can be seen below:

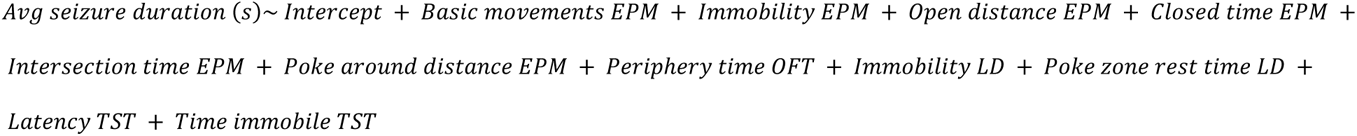

## Results

### Sex dependent effects of ELS on seizure susceptibility

To test the hypothesis that early life stress may worsen epilepsy, we evaluated the impact on seizure susceptibility by recording epileptiform activity in adult male and female ELS and Ctrl mice following acute administration of kainic acid (20 mg/kg KA, *i.p.*). We observed a strong reduction in seizure susceptibility of female *acute-EEG* mice previously exposed to ELS compared to standard-reared controls (Figure 1). In females, ELS significantly lowered the average percent epileptiform activity observed during the 2-hour recording window (Figure 1A) without affecting the latency to the first seizure (Figure 1B). Additionally, all Ctrl (6 of 6, 100.00%) mice entered SE in response to KA administration compared to only one ELS mouse (1 of 8, 11.11%), contributing to the observed increase in latency to SE (Figure 1C). However, we did not observe an impact of ELS on acute seizure-induced elevations in corticosterone (Figure 1D).

**Figure 1:**
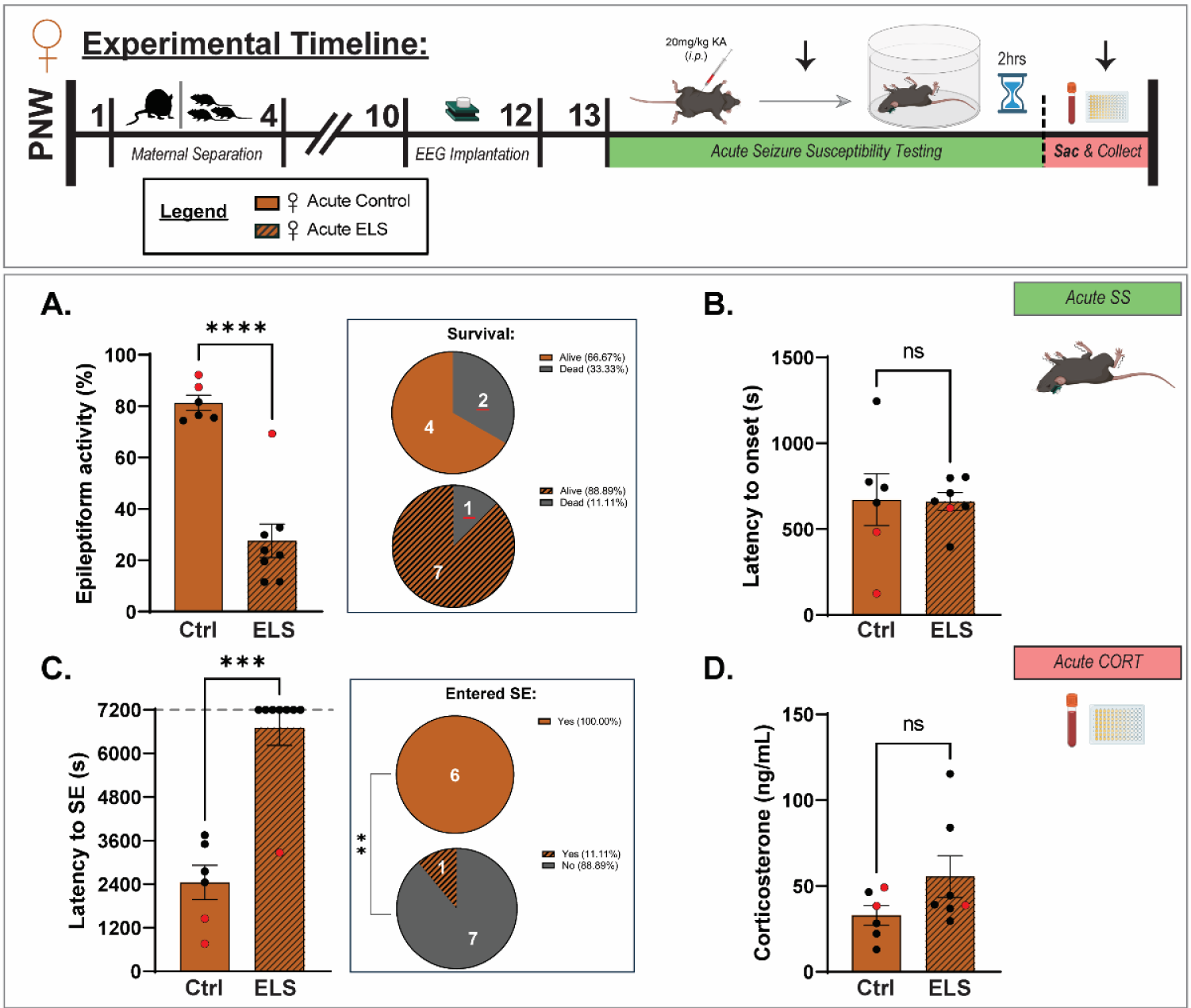
Early life stress reduces seizure susceptibility in female mice. The experimental timeline highlights the timing of procedures for these experiments (top). A, The average percent time exhibiting epileptiform activity in female mice during the two hours post kainic acid administration. B, The average latency to the first ictal event in females following administration of kainic acid. C, The latency to SE in Ctrl and ELS female mice. *Note that animals that did not progress to SE were given the maximum latency. The pie charts depict the number of animals that entered (Yes) or did not enter (No) SE in Ctrl (top) and ELS (bottom). The red dots indicate mice which died prior to the end of the EEG recording. D, The average circulating corticosterone levels measured two hours following kainic acid administration. n = 6-8 mice per experimental group; Student’s unpaired t-test, Mann-Whitney test.

In contrast to female mice, we observed the opposite effect of ELS on acute seizure susceptibility in male mice. Male ELS mice exhibited an increase in the percent time exhibiting epileptiform activity in response to acute kainic acid administration (Figure 2A), with no difference in the latency to seizure onset (Figure 2B). Also, in contrast to females, we did not observe any impact of ELS on the progression to SE (Figure 2C). Neither males nor females demonstrated an impact of ELS on acute seizure-induced elevations in corticosterone (Figures 1D and 2D). These data demonstrate sex dependent impacts of ELS on acute seizure susceptibility with females demonstrating decreased seizure susceptibility while males exhibit an increase in seizure susceptibility.

**Figure 2:**
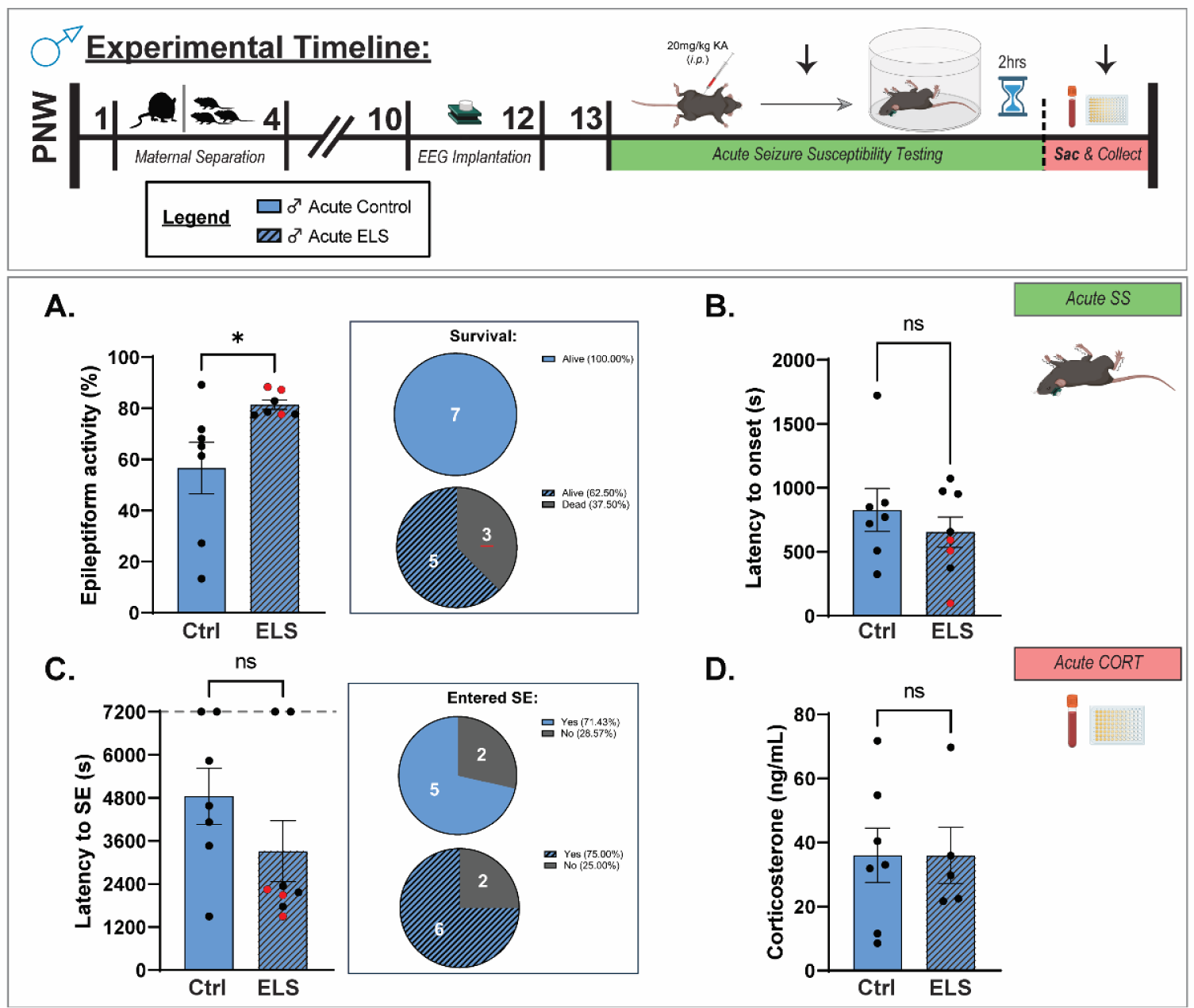
Early life stress increases seizure susceptibility in male mice. The experimental timeline highlights the timing of procedures for these experiments (top). A, The average percent time exhibiting epileptiform activity in male mice during the two hours post kainic acid administration. B, The average latency to the first ictal event in male mice following administration of kainic acid. C, The latency to SE in male Ctrl and ELS mice. *Note that animals that did not progress to SE were given the maximum latency. The pie charts depict the number of animals that entered (Yes) or did not enter (No) SE in Ctrl (top) and ELS (bottom). The red dots indicate mice which died prior to the end of the EEG recording. D, The average circulating corticosterone levels measured two hours following kainic acid administration. n = 5-8 mice per experimental group; Student’s unpaired t-test, Mann-Whitney test.

To further examine the relationship between circulating CORT and acute seizure susceptibility, we performed correlational analyses within *acute-EEG* mice that had both EEG (% epileptiform activity) and CORT (ng/mL) data. As there was no significant effect of sex on acute CORT levels, male and female mice were pooled together to increase sample size. While % epileptiform activity and CORT levels were not significantly correlated in either condition as we have observed in other studies ^2,36^, which may be due to the small n number and skewed distribution in the low versus high seizure activity in the different groups, we did observe that the direction of the relationship was opposite in Ctrl and ELS mice (Figures 3A-B). Further, the seizure-induced elevations in CORT may mask the effects of ELS at higher levels of seizure activity which can only be appreciated in animals with lower seizure frequency.

**Figure 3:**
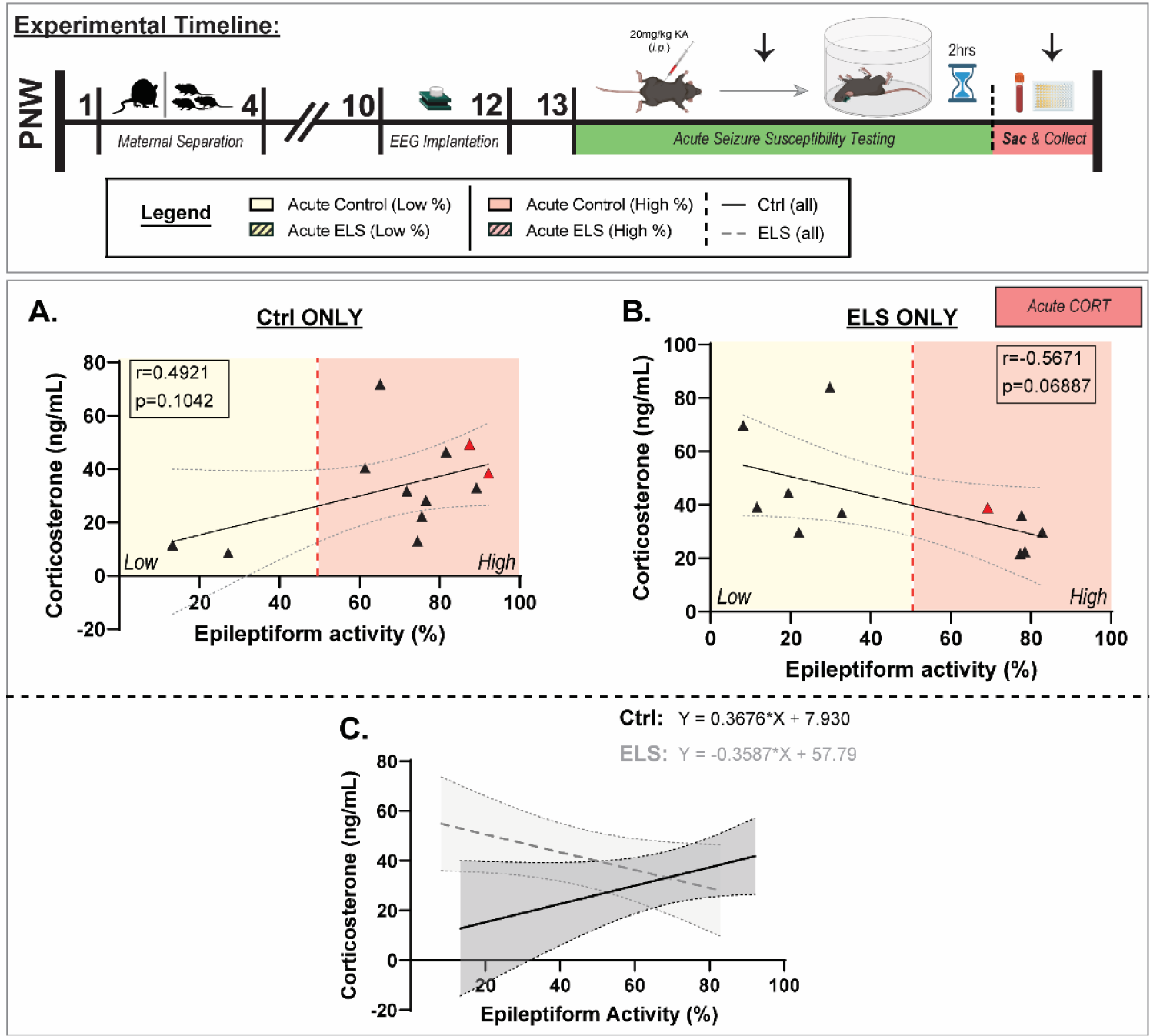
ELS increases seizure-induced corticosterone levels in mice. The experimental timeline highlights the timing of procedures for these experiments (top). The relationship between CORT levels and percent epileptiform activity is shown for Ctrl (A) and ELS (B) mice. C, The overlaid correlations between CORT levels and percent epileptiform activity in Ctrl and ELS mice highlights the differential impact on seizure-induced CORT levels. The red dots indicate mice which died during the EEG recording period. n = 11-12 mice per experimental group; Simple linear regression with Pearson r correlation.

### ELS alters HPA axis function in chronically epileptic female mice which may influence seizure burden

To interrogate the impact of ELS on epilepsy outcomes in a more translationally relevant model, we assessed seizure burden in chronically epileptic ELS and Ctrl mice (*chronic-EEG*) over two weeks of continuous EEG recording. Our data shows that ELS does not alter seizures per hour (Figure 4A, D), seizure duration (Figure B, E), or seizure burden (% time spent seizing) (Figure 4C, F). Although we do not observe an impact of ELS on seizure burden in chronically epileptic mice, we did observe a significant reduction in circulating corticosterone (CORT) levels in chronically epileptic female, but not male mice following ELS (Figure 4G-H) and a positive correlation between CORT levels and epileptiform activity across pooled male and female mice (Figure 4I). These data demonstrate sex differences in the impact of ELS on HPA axis dysfunction in chronically epileptic mice which may influence their seizure burden.

**Figure 4:**
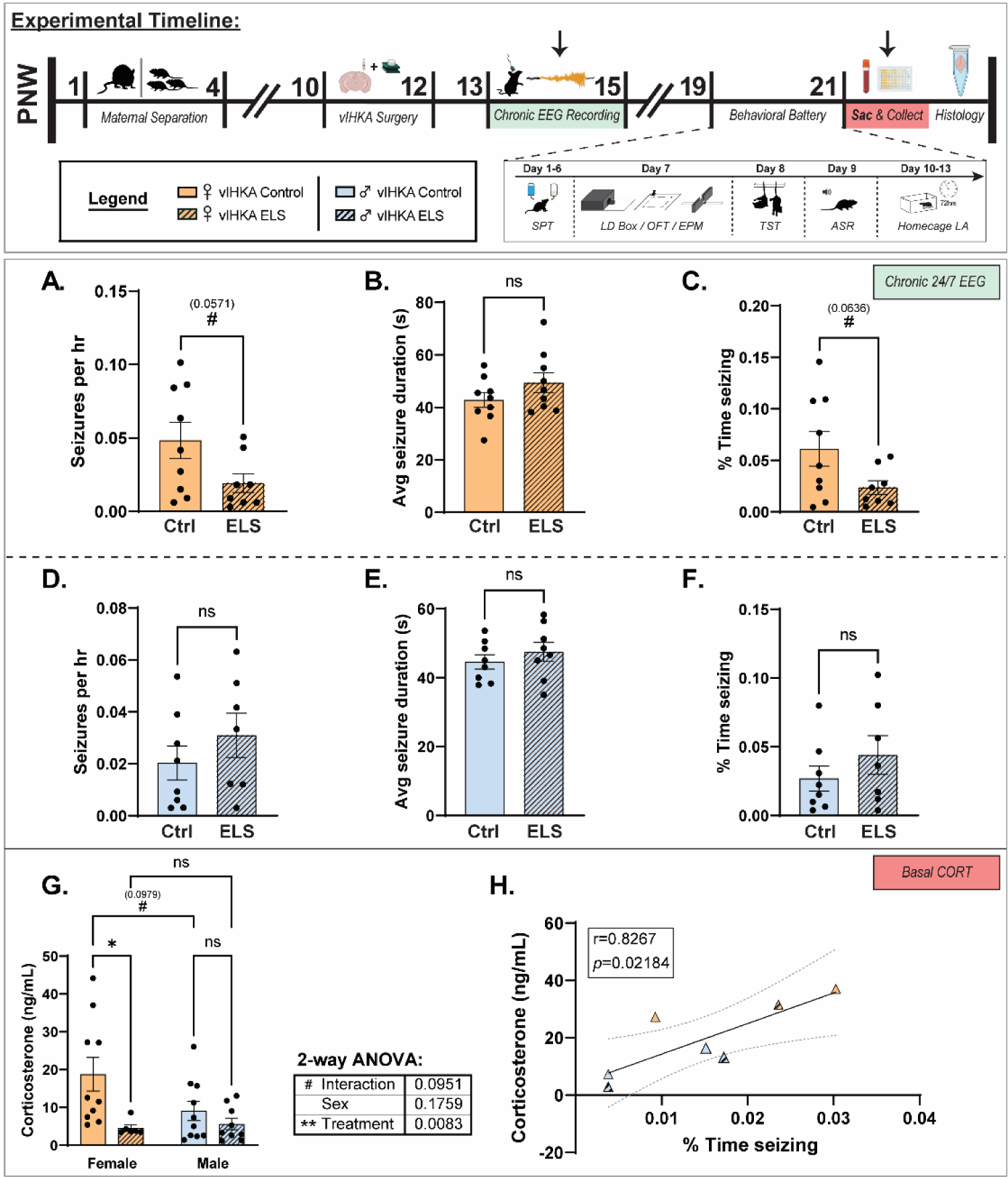
Early life stress impacts HPA axis dysfunction in chronically epileptic mice in a sex dependent manner which may impact seizure burden. The experimental timeline highlights the timing of procedures for these experiments (top). The average number of seizures per hour in chronically epileptic female (A) and male (D) mice. The average seizure duration in chronically epileptic female (B) and male (E) mice. The average seizure burden (% time seizing) in chronically epileptic female (C) and male (F) mice. G, The average corticosterone levels in chronically epileptic Ctrl or ELS female and male mice. H, The relationship between corticosterone levels in chronically epileptic mice and seizure burden (% time seizing). n = 6-10 mice per experimental group; Welch’s t-test, student’s unpaired t-test, 2-way ANOVA, simple linear regression with Pearson r.

### ELS significantly reduces mortality in female chronically epileptic mice

To evaluate the impact of ELS on mortality in chronically epileptic mice, we assessed survivability in the vIHKA model of TLE in adult male and female ELS and Ctrl mice. We observe a significant reduction in mortality associated with chronic epilepsy in female mice subjected to ELS compared to controls (Figure 5A); whereas ELS did not significantly change the survivability of male vIHKA mice compared to controls (Figure 5B). All animals that did not survive were found dead with characteristic hindlimb extension i.e. complete tonic extension (Supplementary Figure S1). To see whether differences in survivability were related to changes in body weight, we measured the weight of mice once prior to the vIHKA surgery and again at the time of sacrifice. We did not observe any significant differences in body weight between vIHKA mice of either sex (Figure 5B, D). Importantly, this characteristic position at death was confirmed as SUDEP in one animal through EEG (Supplementary Figure S1) and across all our EEG animals from our previous publication ^2^. Therefore, we propose that the mortality observed in this mouse model likely occurred due to SUDEP. Based on our previous data demonstrating a role for HPA axis dysfunction in SUDEP risk ^2^, the reduced mortality in females is consistent with the reduced HPA axis dysfunction in females but not males. These data demonstrate for the first time that previous stress exposure may influence SUDEP risk, consistent with our previous findings linking HPA axis dysfunction and SUDEP incidence ^2^.

**Figure 5:**
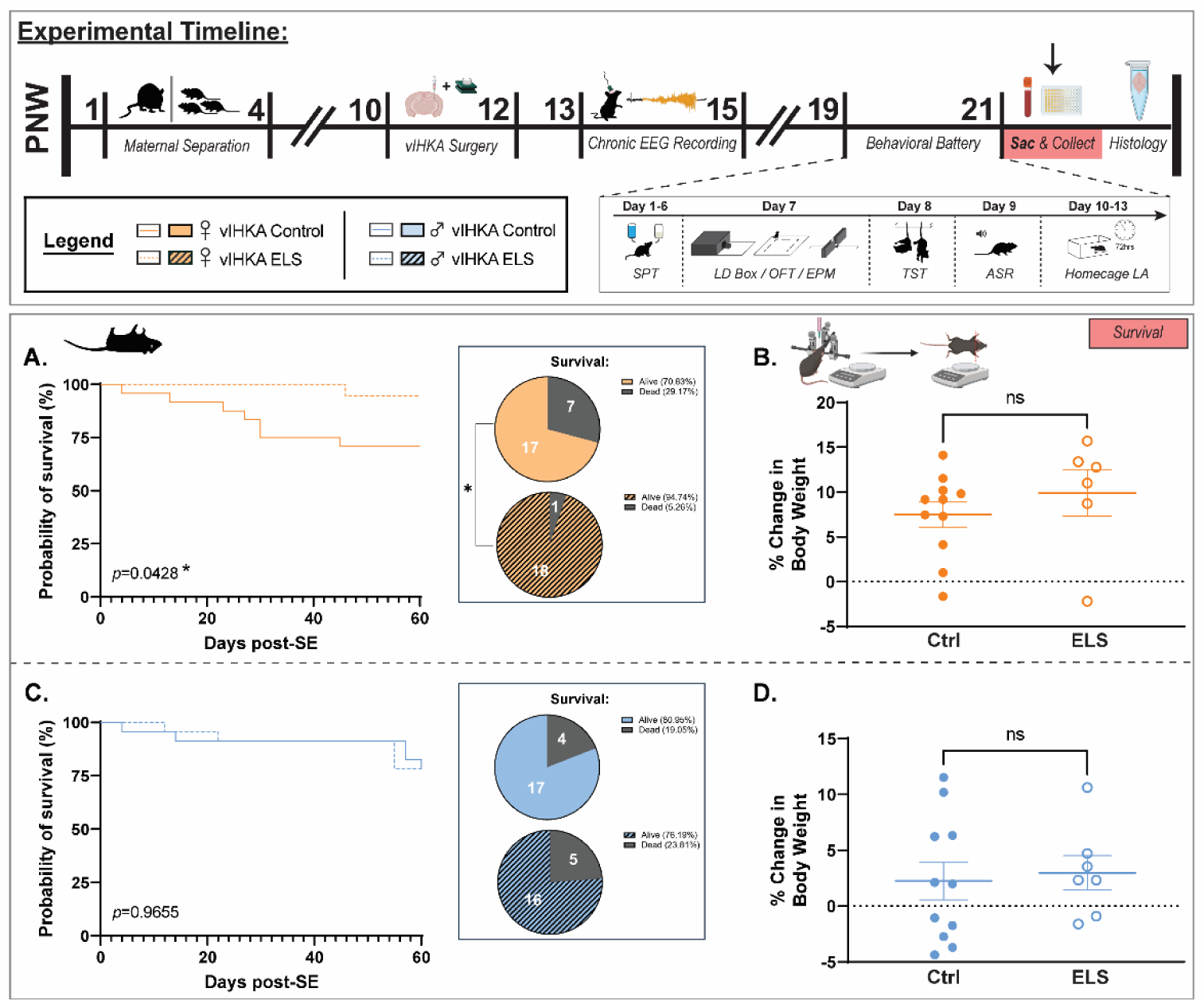
Female chronically epileptic mice subjected to ELS exhibit reduced mortality. The experimental timeline highlights the timing of procedures for these experiments (top). The probability of survival over time is plotted for female (A) and male (C) mice. The pie charts indicate the proportion of mice which survived in Ctrl and ELS female (top) and male (bottom) mice. The average percent change in body weight for chronically epileptic Ctrl and ELS female (B) and male (D) mice; n = 20-24 mice (A, C) or 6-11 mice per experimental group (B, D); Log-rank (Mantel-Cox) test, student’s unpaired t-test.

### ELS influences neuronal activation of the PVN in chronically epileptic mice in a sex dependent manner

To further interrogate the impact of ELS on HPA-axis dysfunction in chronically epileptic mice, we performed c-Fos immunohistochemistry in the paraventricular nucleus of the hypothalamus (PVN), which is at the apex of HPA axis control and has been shown to be activated in response to seizure activity ^3,48^. Interestingly, while we do not observe a significant impact of ELS on c-Fos positive cell count in the PVN in either females (Figures 6A-B) or males (Figures 6A-B), we observe a sex difference in the percentage of c-Fos positive cells in the PVN with male vIHKA mice expressing more c-Fos compared to females (Figure 6B), consistent with the reduced HPA axis activation in chronically epileptic female mice. These findings highlight sex-specific effects of ELS on HPA axis hyperexcitability related to the pathophysiology of epilepsy.

**Figure 6:**
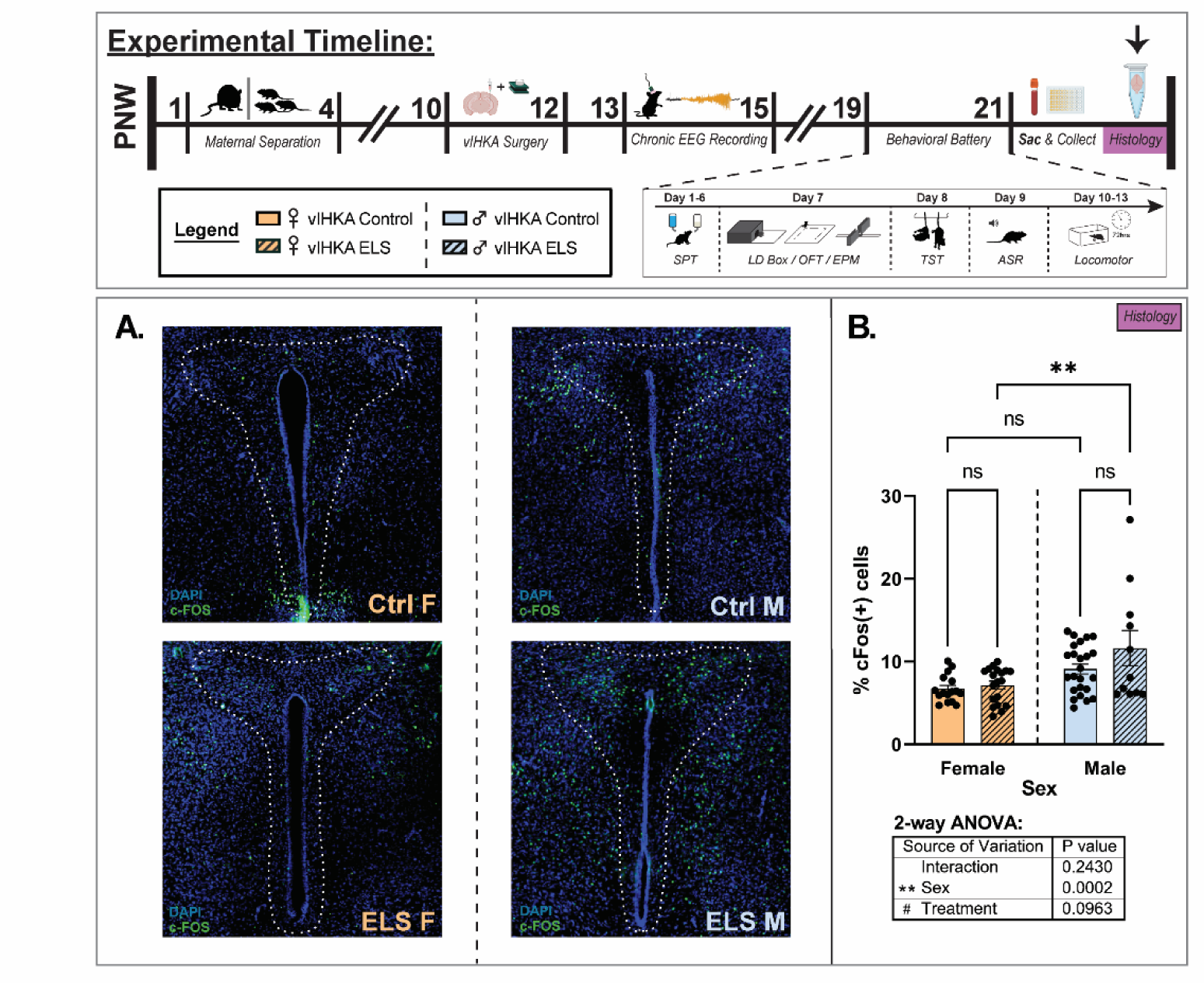
Sex dependent effects of ELS on c-Fos activation in the PVN in chronically epileptic mice. The experimental timeline highlights the timing of procedures for these experiments (top). A, Representative c-Fos immunoreactivity in the PVN of chronically epileptic male and female Ctrl and ELS mice. B, The average percent of c-Fos-positive cells quantified in the PVN of chronically epileptic male and female Ctrl and ELS mice. n = 11-24 sections from 3-5 mice per experimental group; 2-way ANOVA.

### Sex dependent effects of ELS on behavioral outcomes in chronically epileptic mice

Given that HPA axis dysfunction has been implicated in psychiatric comorbidities in epilepsy, we explored the impact of ELS and HPA axis activation on behavioral outcomes in chronically epileptic mice. We assayed avoidance behavior (LD Box, OFT, EPM), the acoustic startle response (ASR), anhedonia (SPT), and general locomotor activity in the home cage (LA). We demonstrate that ELS reduces the max acoustic startle response and increases the distance traveled/exploration in the dark zone of the LD Box (Figure 7B, G). Importantly, ELS did not significantly affect total distance traveled in the LD Box, or in the home cage for female vIHKA mice (Figure 7A, H). No significant differences were observed in behavior between ELS and Ctrl vIHKA mice in the OFT or EPM (7C-E, I-K). Female ELS vIHKA mice also exhibit a decrease in stress-induced helplessness, spending less overall time immobile with a greater latency to immobility during the TST (Figure 8B, C), consistent with the reduced HPA axis dysfunction observed in chronically epileptic female mice. ELS did not significantly affect the average sucrose preference over the 5-day SPT testing period (Figure 8A).

**Figure 7:**
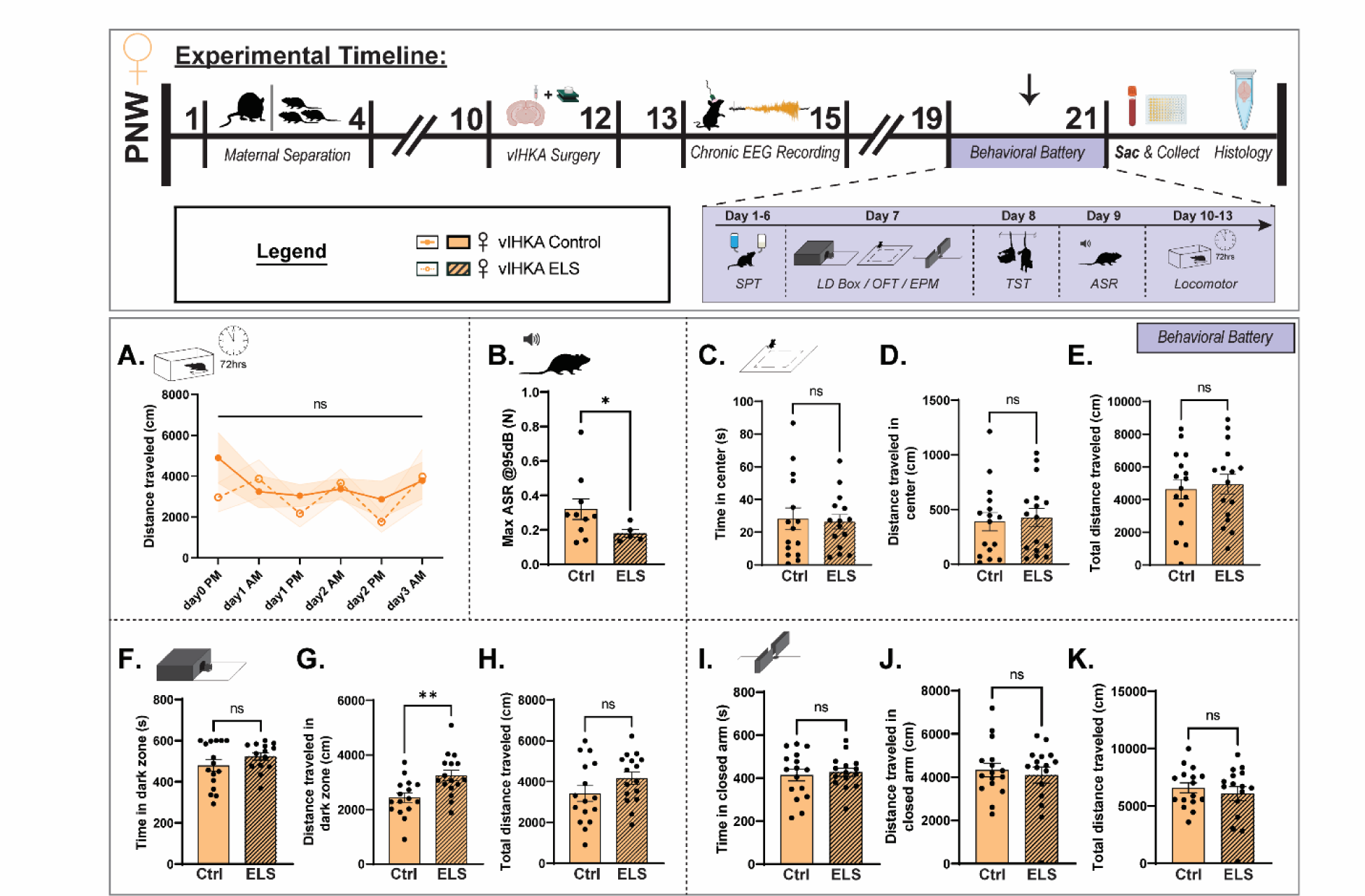
Early life stress induces modest behavioral changes in chronically epileptic female mice. The experimental timeline highlights the timing of procedures for these experiments (top) and the battery of behavioral tests (purple box). A, The average distance traveled in the home cage over days and time. B, The average acoustic startle in Ctrl and ELS chronically epileptic female mice. C-E, The amount of time spent (C) and distance traveled (D) in the center of the open field, with no change in total locomotor activity (E). F-H, The amount of time spent (F) and distance traveled (G) in the dark chamber of the light/dark box, with no change in total locomotor behavior (H). I-K, The amount of time (I) and distance traveled (J) in the open arm of the elevated plus maze, with no difference in total locomotor behavior (K). n = 5-10 mice (A-B) or 14-16 mice (C-K) per experimental group; Mixed-effects analysis, Welch’s t-test, student’s unpaired t-test.

**Figure 8:**
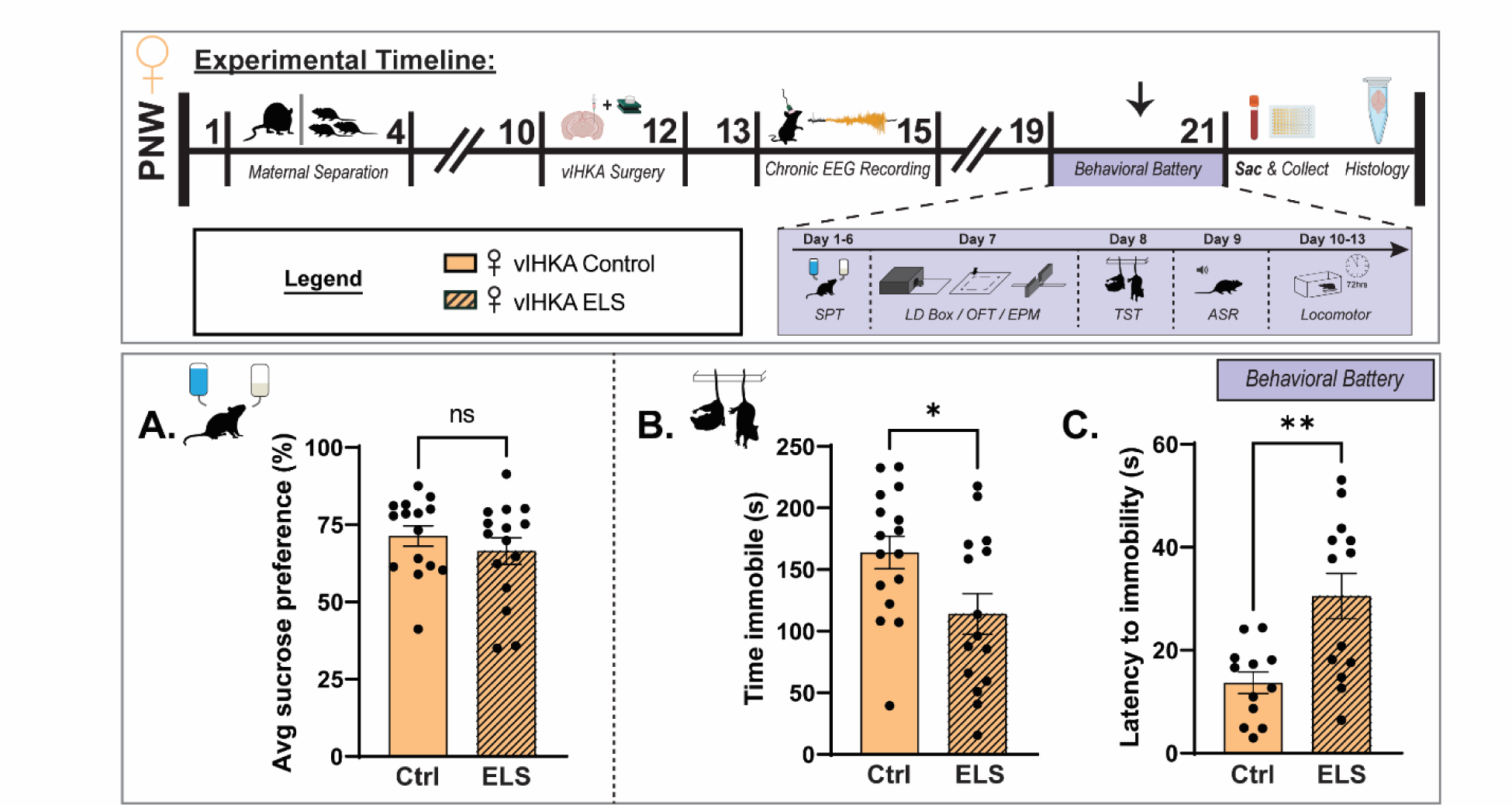
Early life stress reduces stress-induced helplessness behaviors in chronically epileptic female mice. The experimental timeline highlights the timing of procedures for these experiments (top) and the battery of behavioral tests (purple box). A, The sucrose preference in chronically epileptic female Ctrl and ELS mice. B-C, Female chronically epileptic mice subjected to ELS demonstrate a reduced time immobile (B) and an increase in the latency to immobility (C) in the tail suspension test. n = 12-16 mice per experimental group; Student’s unpaired t-test.

While we observed a modest effect of ELS on avoidance and learned-helplessness behaviors in chronically epileptic female mice, this effect was not present in males. Although ELS increased the distance traveled in the center of the OFT and the closed arm of the EPM for male vIHKA mice (Figure 9D, J), it did not significantly affect total distance traveled in either test or in the home cage (Figure 9A, E, K). No significant differences were observed between male ELS and Ctrl vIHKA mice in the LD Box (9F-H), in the time spent in the center of the OFT (Figure 8C), or the closed arms of the EPM (Figure 9I). Neither anhedonic behavior nor stress-induced helplessness behaviors were altered by ELS in male vIHKA mice (Figure 10A-C).

**Figure 9:**
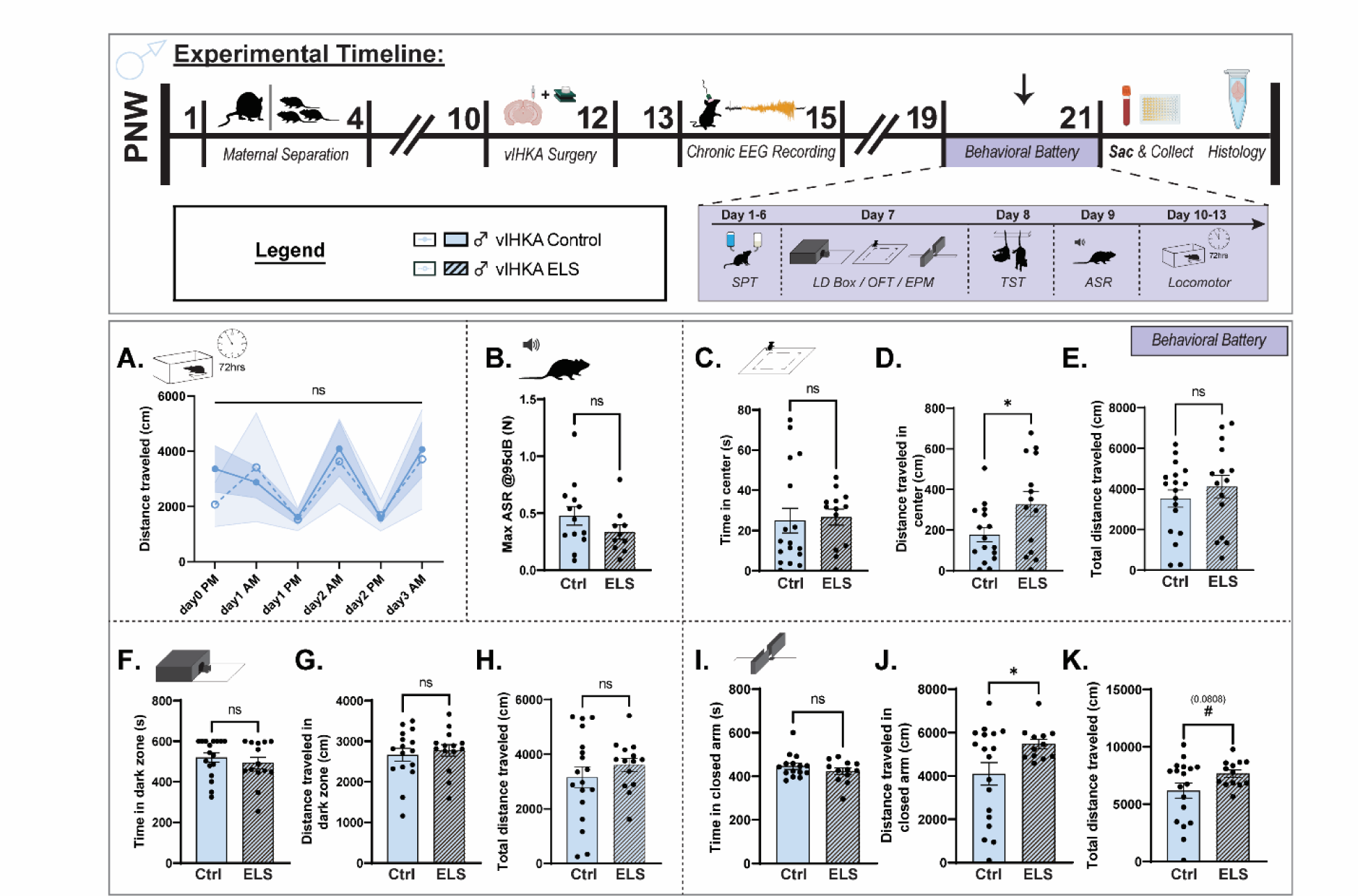
Early life stress does not impact behavioral outcomes in chronically epileptic male mice. The experimental timeline highlights the timing of procedures for these experiments (top) and the battery of behavioral tests (purple box). A, The average distance traveled in the home cage over days and time. B, The average acoustic startle in Ctrl and ELS chronically epileptic male mice. C-E, The amount of time spent (C) and distance traveled (D) in the center of the open field, with no change in total locomotor activity (E). F-H, The amount of time spent (F) and distance traveled (G) in the dark chamber of the light/dark box, with no change in total locomotor behavior (H). I-K, The amount of time (I) and distance traveled (J) in the open arm of the elevated plus maze and total locomotor behavior (K). n = 3-7 mice (A) or 10-18 mice (B-K) per experimental group; Mixed-effects analysis, Welch’s t-test, student’s unpaired t-test.

**Figure 10:**
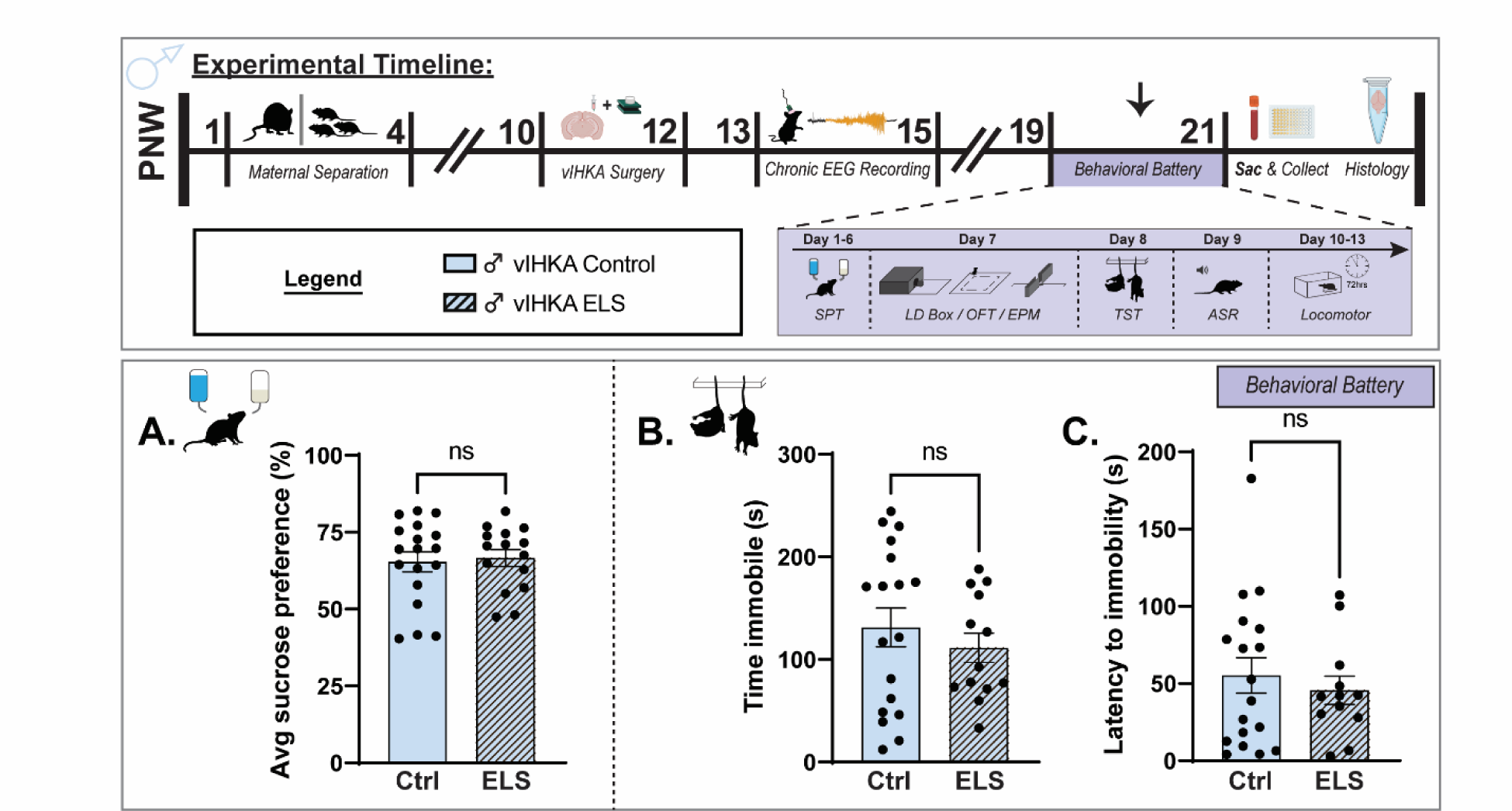
Early life stress does not impact behavioral outcomes in chronically epileptic male mice. The experimental timeline highlights the timing of procedures for these experiments (top) and the battery of behavioral tests (purple box). A, The sucrose preference in chronically epileptic male Ctrl and ELS mice. B-C, The average time spent immobile (B) and the latency to immobility (C) in chronically epileptic male mice in the tail suspension test. n = 12-18 mice per experimental condition; Student’s unpaired t-test.

Taken together, these data highlight sexual dimorphism in altered risk assessment/perceived safety and despair-like behavior of ELS vIHKA mice to further our understanding of how stress experienced in early life impacts vulnerability to comorbid psychiatric illnesses.

Lastly, we assessed the effect of behavioral metrics on predicting seizure outcomes using multi-variable linear regression model. A multiple linear regression analysis was performed to investigate the relationship between behavioral features and the average seizure duration. The overall model was statistically significant (F(11, 10) = 3.673, *p* = 0.0248), accounting for 80.16% of the variance in seizure duration (R² = 0.8016). Among the predictors, ‘Basic Movements EPM’ (β = 2.391, *p* = 0.038), ’Open Distance EPM’ (β = -1.278, *p* = 0.0147), ’Immobility LD’ (β = 1.009, *p* = 0.0009), and ’Latency TST’ (β = 0.8452, *p* = 0.0059) were significant, suggesting associations between these behavioral features and seizure duration.

Multicollinearity was noted, particularly with ’Basic Movements EPM’ (VIF = 50.48) and ’Immobility EPM’ (VIF = 51.58), indicating potential redundancy among these variables. Residuals were normally distributed, as confirmed by multiple normality tests (e.g., Shapiro-Wilk, *p* = 0.7084). Additionally, we looked at the relationship between the same behavioral features and seizure frequency (seizures per hr). In contrast to the model with average seizure duration as the outcome variable, there were no significant associations discovered (F(11, 10) = 0.5322, *p* = 0.8422, R² = 0.3692). These data highlight the associations between behavioral outcomes and seizure metrics in the vIHKA model of TLE with specific relevance to behavioral comorbidities in epilepsy.

## Discussion

It is well accepted that stress is a common seizure trigger. However, the extent to which proximal or previous stress exposures impact epilepsy outcomes are not fully understood. Our work previously demonstrated that seizures activate the HPA axis, contributing to seizure worsening and increasing comorbid behavioral deficits ^3,48^. Further, we demonstrated that HPA axis dysfunction worsens epilepsy outcomes, increasing behavioral deficits associated with chronic epilepsy and increasing SUDEP risk ^2,3^. Here we explore how early life stress impacts epilepsy outcomes, building on previous studies demonstrating sex-dependent effects on seizure outcomes (for review see ^25^). We build on these foundational studies to examine the impact on associated behavioral comorbidities and SUDEP risk.

Our findings demonstrate that ELS impacts seizure susceptibility and epilepsy outcomes in a sex dependent manner. Numerous studies have demonstrated sex differences in the impact of ELS on a range of outcomes ^49^ (for review see ^50^). Previous studies have demonstrated that early life stress impacts seizure outcomes in a sex-dependent manner and has implicated the HPA axis (for review see ^25,34^). We demonstrate a reduction in acute seizure susceptibility in female mice subjected to ELS which is associated with altered seizure-induced activation of the HPA axis. It has been well demonstrated that stress hormones, including corticosterone, are proconvulsant ^6^ and we previously demonstrated that seizure-induced activation of the HPA axis and increased circulating corticosterone levels increases seizure susceptibility ^2,3^. Thus, altered seizure-induced activation of the HPA axis following ELS likely contributes to the impacts of ELS on seizure susceptibility. In fact, we observe a discordant relationship between corticosterone levels and seizure susceptibility following ELS compared to controls.

We previously demonstrated that seizure-induced activation of the HPA axis increased behavioral deficits in chronically epileptic mice ^2,3^. Thus, we examined the impact of ELS, which is associated with blunted HPA axis activation in chronically epileptic mice, on behavioral outcomes. Again, we demonstrate sex differences in the impact of ELS on behavioral outcomes in chronically epileptic mice. We observe that ELS reduces acoustic startle, increased distance traveled/exploration in behavioral tests of avoidance behaviors, and reduced stress-induced helplessness in female mice which is associated with a reduction in corticosterone levels in chronically epileptic mice. These results are consistent with our previous results demonstrating that seizure-induced activation of the HPA axis ^3,48^ and HPA axis dysfunction ^2^ increases vulnerability to behavioral comorbidities in mice with chronic epilepsy. Collectively, these data suggest that HPA axis hyperexcitability increases vulnerability to comorbid behavioral deficits associated with epilepsy, an ELS alters the impact of seizures on HPA axis function which is associated with reduced behavioral comorbidities.

Recently, we demonstrated that HPA axis dysfunction also increases the incidence of SUDEP in chronically epileptic mice and demonstrated a role for neuroendocrine changes in SUDEP risk ^2^. Thus, we evaluated the impact of ELS on mortality in chronically epileptic mice. We demonstrate that female mice subjected to ELS exhibit a reduction in mortality associated with chronic epilepsy. This study demonstrates for the first time that previous stress exposure may influence SUDEP risk. Collectively these studies suggest that HPA axis reactivity and neuroendocrine mediators may contribute to SUDEP risk, requiring further investigation.

Further, it remains to be determined how different types of stressors impact epilepsy outcomes. Our body of work suggests that some chronic stressors such as chronic unpredictable stress induce HPA axis hyperexcitability that worsens epilepsy outcomes ^2,35,36^. Whereas, previous stress exposure which dampens HPA axis function, such as the ELS, paradigm use here can improve epilepsy outcomes.

The exact mechanisms through which altered HPA axis function influences epilepsy outcomes are not fully understood. Stress and prolonged exposure to stress hormones have been shown to alter the structure and function of neurons in the hippocampus (for review see ^51,52^), which is likely to impact epilepsy outcomes particularly in temporal lobe epilepsy. Early life stress has been shown to induce HPA axis reprogramming, altering the neuroendocrine response to stress long-term into adulthood. The ability of ELS to alter HPA axis function, either causing hyper- or hypo-function, is thought to involve the documented impact of ELS on glucocorticoid (GR) and/or mineralocorticoid (MR) signaling and changes in the ability of circulating glucocorticoids to exert negative feedback on the secretion of HPA hormones through binding to GR and MR receptors (for review see ^8^). Further studies are required to determine the exact mechanisms mediating the impact of altered HPA axis function on epilepsy outcomes.

Extensive previous studies demonstrate a link between previous adverse childhood events (ACEs) and the risk for psychiatric illnesses ^53^. Our work suggests that the impact of ACEs on epilepsy outcomes should also be explored, in particular for PWE with comorbid psychiatric illnesses. Stress worsens outcomes across numerous disorders and while stress is a well-accepted seizure trigger, investigation into the impact on the trajectory of epilepsy outcomes has not been prioritized. We advocate for screening PWE for ACEs to inform their vulnerability to psychiatric comorbidities in epilepsy and potentially SUDEP risk.

The data presented here are the first to demonstrate that previous stress exposure influences the trajectory of epilepsy outcomes, altering seizure susceptibility/frequency, associated behavioral deficits, and mortality. We demonstrate that ELS alters seizure-induced HPA axis activation and circulating corticosterone levels which may influence epilepsy outcomes. These data are consistent with previous data from our lab and others ^2–5,7,48,54,55^ and warrant further exploration. Measuring HPA axis function via neuroendocrine markers is straightforward and noninvasive and the emerging evidence suggests that these may be useful biomarkers for predicting the trajectory of epilepsy outcomes.

## Supporting information

Stats Table

## Acknowledgements

Authors would like to thank the members of the Maguire Lab that assisted with mentorship and project guidance. In particular, the authors recognize the contributions of Dr. Kenneth Amaya in the preparation of a Supplement from the NIH which supported this work.

## References

1. Sawyer, N.T., and Escayg, A. (2010). Stress and epilepsy: multiple models, multiple outcomes. J Clin Neurophysiol 27, 445–452. 10.1097/WNP.0b013e3181fe0573.

2. Basu, T., Antonoudiou, P., Weiss, G.L., Coleman, E.M., David, J., Friedman, D., Laze, J., Strain, M.M., Devinsky, O., Boychuk, C.R., and Maguire, J. (2024). Hypothalamic– Pituitary–Adrenal Axis Dysfunction Elevates SUDEP Risk in a Sex-Specific Manner. eneuro *11*, ENEURO.0162–0124.2024. 10.1523/eneuro.0162-24.2024.

3. Hooper, A., Paracha, R., and Maguire, J. (2018). Seizure-induced activation of the HPA axis increases seizure frequency and comorbid depression-like behaviors. Epilepsy Behav 78, 124–133. 10.1016/j.yebeh.2017.10.025.

4. Wulsin, A.C., Herman, J.P., and Danzer, S.C. (2016). RU486 Mitigates Hippocampal Pathology Following Status Epilepticus. Frontiers in Neurology 7, 214. 10.3389/fneur.2016.00214.

5. Wulsin, A.C., Kraus, K.L., Gaitonde, K.D., Suru, V., Arafa, S.R., Packard, B.A., Herman, J.P., and Danzer, S.C. (2021). The glucocorticoid receptor specific modulator CORT108297 reduces brain pathology following status epilepticus. Exp Neurol 341, 113703. 10.1016/j.expneurol.2021.113703.

6. Joëls, M. (2009). Stress, the hippocampus, and epilepsy. Epilepsia 50, 586–597-597. 10.1111/j.1528-1167.2008.01902.x.

7. Wulsin, A.C., Solomon, M.B., Privitera, M.D., Danzer, S.C., and Herman, J.P. (2016). Hypothalamic-pituitary-adrenocortical axis dysfunction in epilepsy. Physiol Behav 166, 22–31. 10.1016/j.physbeh.2016.05.015.

8. Juruena, M.F., Bourne, M., Young, A.H., and Cleare, A.J. (2021). Hypothalamic-Pituitary-Adrenal axis dysfunction by early life stress. Neuroscience Letters 759, 136037. 10.1016/j.neulet.2021.136037.

9. Lupien, S.J., McEwen, B.S., Gunnar, M.R., and Heim, C. (2009). Effects of stress throughout the lifespan on the brain, behaviour and cognition. Nature Reviews Neuroscience 10, 434–445. 10.1038/nrn2639.

10. Andersen, S.L. (2003). Trajectories of brain development: point of vulnerability or window of opportunity? Neuroscience & Biobehavioral Reviews 27, 3–18. 10.1016/S0149-7634(03)00005-8.

11. 11. Schalkwijk, F., Van Someren, E.J.W., Nicolai, N.J., Uijttewaal, J.L., and Wassing, R. (2023). From childhood trauma to hyperarousal in adults: The mediating effect of maladaptive shame coping and insomnia. Frontiers in human neuroscience 17, 990581. 10.3389/fnhum.2023.990581.

12. Pollak, S.D., and Sinha, P. (2002). Effects of early experience on children’s recognition of facial displays of emotion. Developmental psychology 38, 784–791. 10.1037//0012-1649.38.5.784.

13. Pollak, S.D., and Tolley-Schell, S.A. (2003). Selective attention to facial emotion in physically abused children. Journal of abnormal psychology 112, 323–338. 10.1037/0021-843x.112.3.323.

14. Masson, M., East-Richard, C., and Cellard, C. (2016). A meta-analysis on the impact of psychiatric disorders and maltreatment on cognition. Neuropsychology 30, 143–156. 10.1037/neu0000228.

15. Geoffroy, M.C., Pinto Pereira, S., Li, L., and Power, C. (2016). Child Neglect and Maltreatment and Childhood-to-Adulthood Cognition and Mental Health in a Prospective Birth Cohort. Journal of the American Academy of Child and Adolescent Psychiatry 55, 33–40.e33. 10.1016/j.jaac.2015.10.012.

16. Chen, Y., and Baram, T.Z. (2016). Toward Understanding How Early-Life Stress Reprograms Cognitive and Emotional Brain Networks. Neuropsychopharmacology 41, 197–206. 10.1038/npp.2015.181.

17. Walker, C.-D., Bath, K.G., Joels, M., Korosi, A., Larauche, M., Lucassen, P.J., Morris, M.J., Raineki, C., Roth, T.L., Sullivan, R.M., et al. (2017). Chronic early life stress induced by limited bedding and nesting (LBN) material in rodents: critical considerations of methodology, outcomes and translational potential. Stress 20, 421–448. 10.1080/10253890.2017.1343296.

18. Schmidt, M.V., Wang, X.-D., and Meijer, O.C. (2011). Early life stress paradigms in rodents: potential animal models of depression? Psychopharmacology 214, 131–140. 10.1007/s00213-010-2096-0.

19. Elzinga, B.M., Roelofs, K., Tollenaar, M.S., Bakvis, P., van Pelt, J., and Spinhoven, P. (2008). Diminished cortisol responses to psychosocial stress associated with lifetime adverse events a study among healthy young subjects. Psychoneuroendocrinology 33, 227–237. 10.1016/j.psyneuen.2007.11.004.

20. Yang, J.Z., Kang, C.Y., Yuan, J., Zhang, Y., Wei, Y.J., Xu, L., Zhou, F., and Fan, X. (2021). Effect of adverse childhood experiences on hypothalamic-pituitary-adrenal (HPA) axis function and antidepressant efficacy in untreated first episode patients with major depressive disorder. Psychoneuroendocrinology 134, 105432. 10.1016/j.psyneuen.2021.105432.

21. Murphy, F., Nasa, A., Cullinane, D., Raajakesary, K., Gazzaz, A., Sooknarine, V., Haines, M., Roman, E., Kelly, L., O’Neill, A., et al. (2022). Childhood Trauma, the HPA Axis and Psychiatric Illnesses: A Targeted Literature Synthesis. Frontiers in Psychiatry 13. 10.3389/fpsyt.2022.748372.

22. Gluckman, P.D., Hanson, M.A., and Beedle, A.S. (2007). Early life events and their consequences for later disease: a life history and evolutionary perspective. American journal of human biology : the official journal of the Human Biology Council 19, 1–19. 10.1002/ajhb.20590.

23. 23. van Bodegom, M., Homberg, J.R., and Henckens, M. (2017). Modulation of the Hypothalamic-Pituitary-Adrenal Axis by Early Life Stress Exposure. Front Cell Neurosci 11, 87. 10.3389/fncel.2017.00087.

24. 24. van Campen, J.S., Jansen, F.E., de Graan, P.N.E., Braun, K.P.J., and Joels, M. (2014). Early life stress in epilepsy: A seizure precipitant and risk factor for epileptogenesis. Epilepsy & Behavior 38, 160–171. 10.1016/j.yebeh.2013.09.029.

25. Jones, N.C., O’Brien, T.J., and Carmant, L. (2014). Interaction between sex and early-life stress: Influence on epileptogenesis and epilepsy comorbidities. Neurobiology of Disease 72, 233–241. 10.1016/j.nbd.2014.09.004.

26. Koe, A.S., Jones, N.C., and Salzberg, M.R. (2009). Early life stress as an influence on limbic epilepsy: an hypothesis whose time has come? Front Behav Neurosci 3, 24. 10.3389/neuro.08.024.2009.

27. Frye, C.A., and Bayon, L.E. (1999). Prenatal stress reduces the effectiveness of the neurosteroid 3α, 5α-THP to block kainic-acid-induced seizures. Developmental Psychobiology: The Journal of the International Society for Developmental Psychobiology 34, 227–234.

28. Edwards, H.E., Dortok, D., Tam, J., Won, D., and Burnham, W.M. (2002). Prenatal stress alters seizure thresholds and the development of kindled seizures in infant and adult rats. Hormones and behavior 42, 437–447.

29. Sadaghiani, M.M., and Saboory, E. (2010). Prenatal stress potentiates pilocarpine-induced epileptic behaviors in infant rats both time and sex dependently. Epilepsy & Behavior 18, 166–170.

30. Desgent, S., Duss, S., Sanon, N.T., Lema, P., Lévesque, M., Hébert, D., Rébillard, R.-M., Bibeau, K., Brochu, M., and Carmant, L. (2012). Early-life stress is associated with gender-based vulnerability to epileptogenesis in rat pups.

31. Kumar, G., Jones, N.C., Morris, M.J., Rees, S., O’Brien, T.J., and Salzberg, M.R. (2011). Early life stress enhancement of limbic epileptogenesis in adult rats: mechanistic insights. PLoS One 6, e24033.

32. Salzberg, M., Kumar, G., Supit, L., Jones, N.C., Morris, M.J., Rees, S., and O’Brien, T.J. (2007). Early postnatal stress confers enduring vulnerability to limbic epileptogenesis. Epilepsia 48, 2079–2085.

33. Rupasinghe, R., Dezsi, G., Ozturk, E., Carron, S., Hudson, M.R., Casillas-Espinosa, P.M., and Jones, N.C. (2022). Early life adversity accelerates epileptogenesis and enhances depression-like behaviors in rats. Experimental Neurology 354, 114088. 10.1016/j.expneurol.2022.114088.

34. Koe, A.S., Salzberg, M.R., Morris, M.J., O’Brien, T.J., and Jones, N.C. (2014). Early life maternal separation stress augmentation of limbic epileptogenesis: The role of corticosterone and HPA axis programming. Psychoneuroendocrinology 42, 124–133. 10.1016/j.psyneuen.2014.01.009.

35. Hooper, A., Paracha, R., and Maguire, J. (2017). Seizure-induced activation of the HPA axis increases seizure frequency and comorbid depression-like behaviors. Epilepsy & Behavior 78, 124–133.

36. O’Toole, K.K., Hooper, A., Wakefield, S., and Maguire, J. (2013). Seizure-induced disinhibition of the HPA axis increases seizure susceptibility. Epilepsy Research 108(*1*), 29–43.

37. Wulsin, A.C., Herman, J.P., and Danzer, S.C. (2016). RU486 Mitigates Hippocampal Pathology Following Status Epilepticus. Front Neurol 7, 214.

38. Salpekar, J.A., Basu, T., Thangaraj, S., and Maguire, J. (2020). The intersections of stress, anxiety and epilepsy. In International Review of Neurobiology, (Academic Press). 10.1016/bs.irn.2020.02.001.

39. Stone, B.T., Antonoudiou, P., Teboul, E., Scarpa, G., Weiss, G., and Maguire, J.L. (2023). Early life stress impairs VTA coordination of BLA network and behavioral states. bioRxiv, 2023.2009.2016.558081. 10.1101/2023.09.16.558081.

40. 40. Scarpa, G., Antonoudiou, P., Weiss, G., Stone, B., and Maguire, J.L. (2024). Sex-dependent effects of early life stress on network and behavioral states. bioRxiv, 2024.2005.2010.593547. 10.1101/2024.05.10.593547.

41. Trinka, E., Cock, H., Hesdorffer, D., Rossetti, A.O., Scheffer, I.E., Shinnar, S., Shorvon, S., and Lowenstein, D.H. (2015). A definition and classification of status epilepticus--Report of the ILAE Task Force on Classification of Status Epilepticus. Epilepsia 56, 1515–1523. 10.1111/epi.13121.

42. Lowenstein, D.H., Bleck, T., and Macdonald, R.L. (1999). It’s time to revise the definition of status epilepticus. Epilepsia 40, 120–122. 10.1111/j.1528-1157.1999.tb02000.x.

43. Zeidler, Z., Brandt-Fontaine, M., Leintz, C., Krook-Magnuson, C., Netoff, T., and Krook-Magnuson, E. (2018). Targeting the Mouse Ventral Hippocampus in the Intrahippocampal Kainic Acid Model of Temporal Lobe Epilepsy. eNeuro 5. 10.1523/ENEURO.0158-18.2018.

44. Antonoudiou, P., Basu, T., and Maguire, J. (2023). Semi-automated seizure detection using interpretable machine learning models. bioRxiv, 2023.2010.2025.563903. 10.1101/2023.10.25.563903.

45. Melon, L.C., Hooper, A., Yang, X., Moss, S.J., and Maguire, J. (2018). Inability to suppress the stress-induced activation of the HPA axis during the peripartum period engenders deficits in postpartum behaviors in mice. Psychoneuroendocrinology 90, 182–193. 10.1016/j.psyneuen.2017.12.003.

46. Walton, N.L., Antonoudiou, P., Barros, L., Dargan, T., DiLeo, A., Evans-Strong, A., Gabby, J., Howard, S., Paracha, R., Sánchez, E.J., et al. (2023). Impaired Endogenous Neurosteroid Signaling Contributes to Behavioral Deficits Associated With Chronic Stress. Biol Psychiatry 94, 249–261. 10.1016/j.biopsych.2023.01.022.

47. Antonoudiou, P., Colmers, P.L.W., Walton, N.L., Weiss, G.L., Smith, A.C., Nguyen, D.P., Lewis, M., Quirk, M.C., Barros, L., Melon, L.C., and Maguire, J.L. (2022). Allopregnanolone Mediates Affective Switching Through Modulation of Oscillatory States in the Basolateral Amygdala. Biol Psychiatry 91, 283–293. 10.1016/j.biopsych.2021.07.017.

48. O’Toole, K.K., Hooper, A., Wakefield, S., and Maguire, J. (2014). Seizure-induced disinhibition of the HPA axis increases seizure susceptibility. Epilepsy research 108, 29–43. 10.1016/j.eplepsyres.2013.10.013.

49. Goodwill, H.L., Manzano-Nieves, G., Gallo, M., Lee, H.-I., Oyerinde, E., Serre, T., and Bath, K.G. (2019). Early life stress leads to sex differences in development of depressive-like outcomes in a mouse model. Neuropsychopharmacology 44, 711–720. 10.1038/s41386-018-0195-5.

50. Bale, T.L., and Epperson, C.N. (2015). Sex differences and stress across the lifespan. Nature Neuroscience 18, 1413–1420. 10.1038/nn.4112.

51. McEwen, B.S., Nasca, C., and Gray, J.D. (2016). Stress Effects on Neuronal Structure: Hippocampus, Amygdala, and Prefrontal Cortex. Neuropsychopharmacology 41, 3–23. 10.1038/npp.2015.171.

52. Joels, M. (2009). Stress, the hippocampus, and epilepsy. Epilepsia 50, 586–597.

53. Daníelsdóttir, H.B., Aspelund, T., Shen, Q., Halldorsdottir, T., Jakobsdóttir, J., Song, H., Lu, D., Kuja-Halkola, R., Larsson, H., Fall, K., et al. (2024). Adverse Childhood Experiences and Adult Mental Health Outcomes. JAMA Psychiatry 81, 586–594. 10.1001/jamapsychiatry.2024.0039.

54. Wulsin, A.C., Franco-Villanueva, A., Romancheck, C., Morano, R.L., Smith, B.L., Packard, B.A., Danzer, S.C., and Herman, J.P. (2018). Functional disruption of stress modulatory circuits in a model of temporal lobe epilepsy. PLOS ONE 13, e0197955. 10.1371/journal.pone.0197955.

55. Basu, T., Maguire, J., and Salpekar, J.A. (2021). Hypothalamic-pituitary-adrenal axis targets for the treatment of epilepsy. Neurosci Lett 746, 135618. 10.1016/j.neulet.2020.135618.

