## Supplemental Material for "Early life stress influences epilepsy outcomes in mice"

**Extended Data/Supplemental Information**

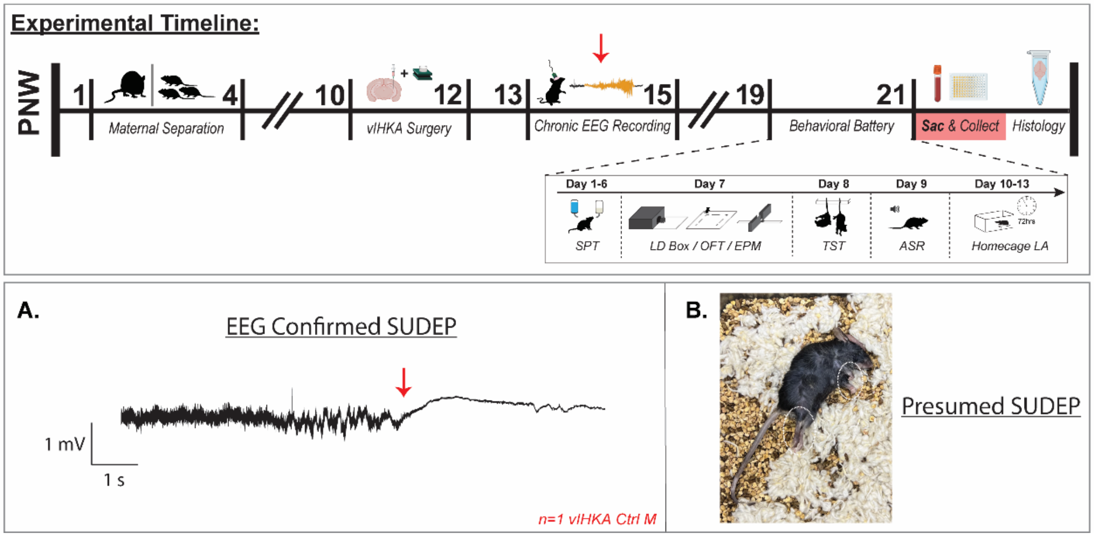


**Figure S1: Representative EEG trace and photo of confirmed and presumed SUDEP.** The experimental timeline highlights the timing of EEG confirmed SUDEP (red arrow) in relation to the study, whereas presumed SUDEP occurred at various timepoints during PNW 13-21. A, Representative trace showing the only case of confirmed SUDEP from one male vIHKA Ctrl mouse. B, All animals that did not survive were found in complete tonic extension, including the one Ctrl mouse that had EEG confirmed SUDEP.

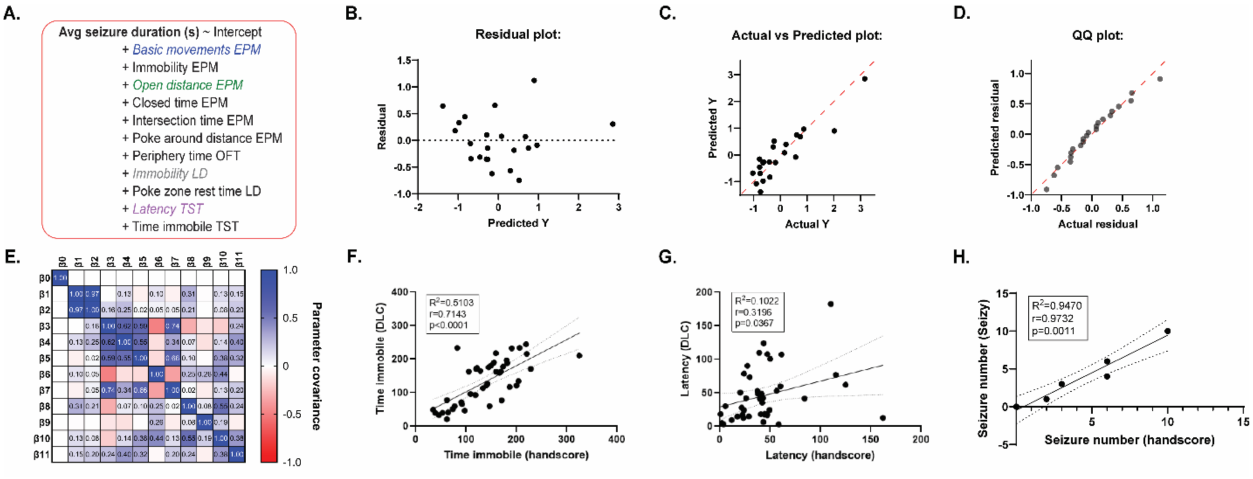


**Figure S2: Multiple Linear Regression analysis and automated scoring validation.** A, The equation used for multiple linear regression analysis performed on normalized data, with significant independent variables ($p<0.05$) in color text and italicized. B-D, The residuals of the multiple linear regression analysis plotted in three different ways: predicted y-value vs. residual (B), actual y-value vs. predicted y-value (C), and the actual residual vs. the predicted residual (D). E, Correlation matrix showing the covariance between the 12 predicted parameters (β0-11) in the multiple linear regression analysis. F-G, The relationship between hand-scored videos of the tail suspension test and the same videos scored using DeepLabCut (DLC) across the two reported metrics: total time spent immobile (F) and the latency to immobility (G). H, The relationship between the number of seizures detected through hand-scored EEG recordings from chronically epileptic mice and the number of seizures detected by our automated seizure detection app Seizy in the same EEG recordings. n = 22 mice (A-E) or 43 videos (F-G) or 6 24hr recordings (H); Multiple linear regression, simple linear regression with Pearson r.

| Analysis of Variance | SS | DF | MS | F (DFn, DFd) | Estimate | Standard error | P value | VIF | R² |
|---|---|---|---|---|---|---|---|---|---|
| Regression (β0) | 17.64 | 11 | 1.603 | F (11, 10) = 3.673 | -7.69E-16 | 0.1408 | P=0.0248 | | |
| **Basic movements EPM (β1)** | **2.492** | **1** | **2.492** | **F (1, 10) = 5.709** | 2.391 | 1.001 | **P=0.0380** | 50.48 | 0.9802 |
| Immobility EPM (β2) | 0.8429 | 1 | 0.8429 | F (1, 10) = 1.931 | 1.406 | 1.012 | P=0.1948 | 51.58 | 0.9806 |
| **Open distance EPM (β3)** | **3.78** | **1** | **3.78** | **F (1, 10) = 8.661** | -1.278 | 0.4343 | **P=0.0147** | 9.508 | 0.8948 |
| Closed time EPM (β4) | 1.31 | 1 | 1.31 | F (1, 10) = 3.001 | -0.4288 | 0.2475 | P=0.1139 | 3.087 | 0.6761 |
| Intersection time EPM (β5) | 0.04603 | 1 | 0.04603 | F (1, 10) = 0.1055 | 0.07836 | 0.2413 | P=0.7521 | 2.934 | 0.6592 |
| Poke around distance EPM (β6) | 0.2667 | 1 | 0.2667 | F (1, 10) = 0.6110 | 0.1482 | 0.1896 | P=0.4525 | 1.813 | 0.4484 |
| Periphery time OFT (β7) | 0.8285 | 1 | 0.8285 | F (1, 10) = 1.898 | -0.4138 | 0.3003 | P=0.1983 | 4.546 | 0.78 |
| **Immobility LD (β8)** | **9.414** | **1** | **9.414** | **F (1, 10) = 21.57** | 1.009 | 0.2172 | **P=0.0009** | 2.378 | 0.5795 |
| Poke zone rest time LD (β9) | 0.04814 | 1 | 0.04814 | F (1, 10) = 0.1103 | -0.05128 | 0.1544 | P=0.7467 | 1.202 | 0.1678 |
| **Latency TST (β10)** | **5.288** | **1** | **5.288** | **F (1, 10) = 12.12** | 0.8452 | 0.2428 | **P=0.0059** | 2.972 | 0.6635 |
| Time immobile TST (β11) | 0.2729 | 1 | 0.2729 | F (1, 10) = 0.6254 | 0.1391 | 0.1759 | P=0.4474 | 1.56 | 0.3589 |
| Residual | 4.364 | 10 | 0.4364 | | | | | | |
| Total | 22 | 21 | | | | | | | |

**Table S1: Multiple Linear Regression results.**

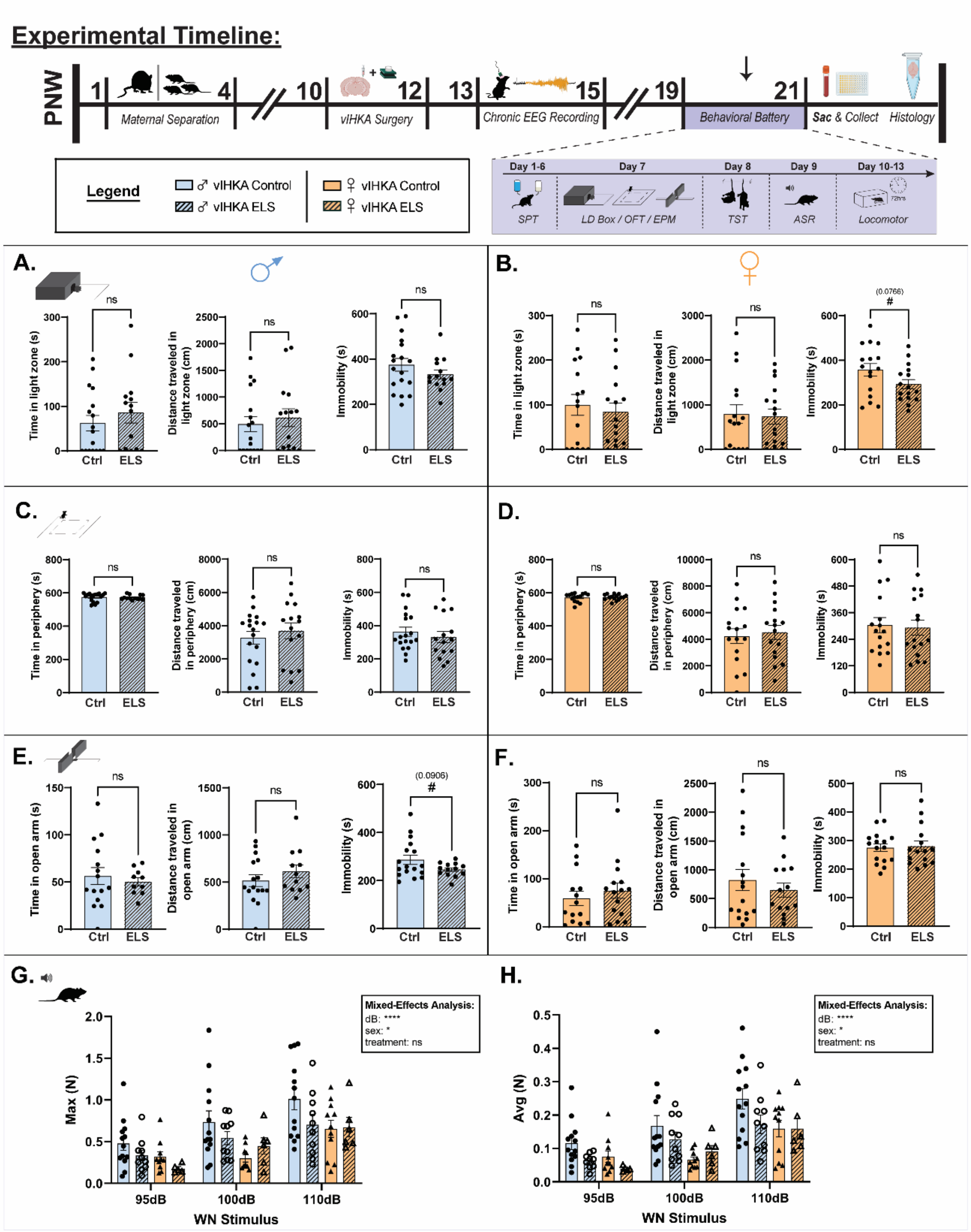


**Figure S3: Extended data from the battery of behavioral tests performed on chronically epileptic Ctrl and ELS mice.** The experimental timeline highlights the timing of procedures for these experiments (top) and the battery of behavioral tests (purple box). A-B, The amount of time spent and distance traveled in the light zone of the light/dark box, as well as the total time spent immobile in Ctrl and ELS chronically epileptic male (A) and female (B) mice. C-D, The amount of time spent and distance traveled in periphery of the open field test, as well as the total time spent immobile in Ctrl and ELS chronically epileptic male (C) and female (D) mice. E-F,

The amount of time spent and distance traveled in the open arms of the elevated plus maze, as well as the total time spent immobile in Ctrl and ELS chronically epileptic male (E) and female (F) mice. G-H, The maximum (G) and average (H) acoustic startle response across three decibels of stimulus presentation in Ctrl and ELS chronically epileptic male and female mice. n = 10-18 mice (A-F) or 6-13 mice (G-H) per experimental group; Student's unpaired t-test, mixed-effects model (REML).

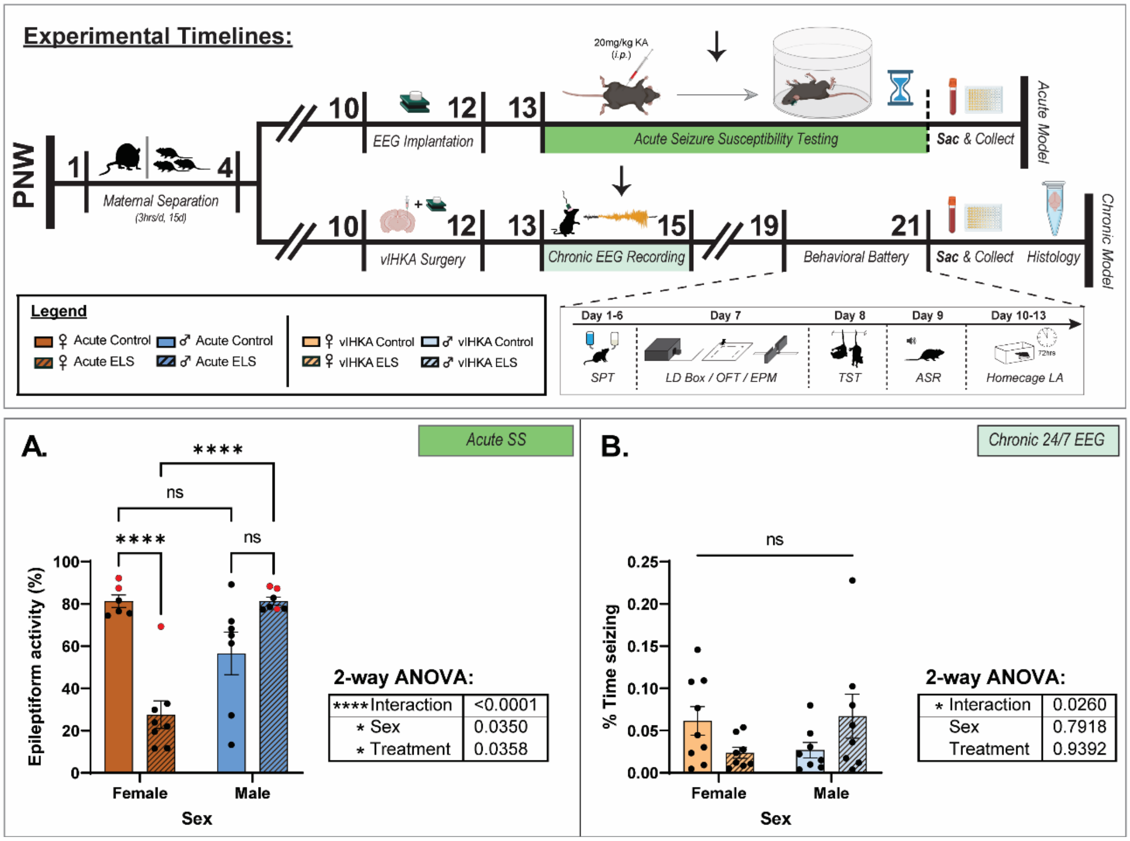


**Figure S4: Comparisons across multiple conditions reveal sex differences in the acute model of seizure susceptibility, but not the chronic model.** The experimental timeline highlights the timing of procedures for these experiments in the acute model (top) and the chronic model (bottom). A, The average percent time exhibiting epileptiform activity Ctrl and ELS mice of both sexes during the two hours following kainic acid administration. The red dots indicate mice which died prior to the end of the EEG recording. B, The average seizure burden (% time seizing) in chronically epileptic Ctrl and ELS mice of both sexes. n = 6-9 mice per experimental group; 2-way ANOVA.
