## Supplementary material for "Early life stress influences epilepsy outcomes in mice": Stats Table

**Statistics Tables:**

**Figure 1- Acute SS (Female)**

| Description | Figure | Statistical test | Results |
|---|---|---|---|
| Epileptiform activity (%): Acute Ctrl F v Acute ELS F | 1A | Unpaired t test | t=6.699, df=12, p<0.0001 |
| Latency to onset: Acute Ctrl F vs. Acu ELS F | 1B | Unpaired t test | t=0.06941, df=11, p=0.9459 |
| Latency to SE: Acute Ctrl F vs. Acute F | 1C | Mann-Whitney | U=2, p=0.0010 |
| Entered SE (%): Acute Ctrl F vs. Acut ELS F | 1C | Mann-Whitney | U=3, p=0.0047 |
| CORT: Acute Ctrl F vs. Acute ELS F | 1D | Unpaired t test | t=1.599, df=11, p=0.1381 |

**Figure 2- Acute SS (Male)**

| Description | Figure | Statistical test | Results |
|---|---|---|---|
| Epileptiform activity (%): Acute Ctrl M Acute ELS M | 2A | Unpaired t test | t=2.422, df=12, p=0.0322 |
| Latency to onset: Acute Ctrl M vs. Acu ELS M | 2B | Unpaired t test | t=0.8603, df=13, p=0.405 |
| Latency to SE: Acute Ctrl M vs. Acute M | 2C | Mann-Whitney | U=18, p=0.2600 |
| Entered SE (%): Acute Ctrl M vs. Acu ELS M | 2C | Mann-Whitney | U=27, p>0.9999 |
| CORT: Acute Ctrl M vs. Acute ELS M | 2D | Unpaired t test | t=0.007683, df=10, p=0.9940 |

**Figure 3- Acute EEG + CORT**

| Description | Figure | Statistical test | Results |
|---|---|---|---|
| Acute CORT vs. Epileptiform activity ( CTRL only | 3A | Simple linear regression | Y = 0.3676*X + 7.930 |
| Acute CORT vs. Epileptiform activity ( CTRL only | 3A | Pearson r correlation | r=0.4921, $R^2$=0.2421, p=0.1042 |
| Acute CORT vs. Epileptiform activity ( ELS only | 3B | Simple linear regression | Y = -0.3587*X + 57.79 |
| Acute CORT vs. Epileptiform activity ( ELS only | 3B | Pearson r correlation | r=-0.5671, $R^2$=0.3216, p=0.0689 |

**Figure 4- Chronic EEG + CORT**

| Description | Figure | Statistical test | Results |
|---|---|---|---|
| Seizures per hr: vIHKA Ctrl F vs. vIHKA ELS F | 4A | Welch's t test | t=2.105, df=11.97, p=0.057 |

| | | | |
|---|---|---|---|
| Avg seizure duration: vIHKA Ctrl F vIHKA ELS F | 4B | Unpaired t test | t=1.389, df=16, p=0.1837 |
| % time seizing: vIHKA Ctrl F vs. vIHKA ELS F | 4C | Welch's t test | t=2.077, df=10.36, p=0.0636 |
| Seizures per hr: vIHKA Ctrl M vs. vIHKA ELS M | 4D | Unpaired t test | t=0.9954, df=13, p=0.3377 |
| Avg seizure duration: vIHKA Ctrl M vs. vIHKA ELS M | 4E | Unpaired t test | t=0.8368, df=14, p=0.4168 |
| % time seizing: vIHKA Ctrl M vs. vIHKA ELS M | 4F | Unpaired t test | t=1.050, df=13, p=0.3126 |
| Basal CORT | 4G | 2-Way ANOVA | Interaction: F (1, 31) = 2.964, p=0.0951 |
| Basal CORT | 4G | 2-Way ANOVA | Sex: F (1, 31) = 1.919, p=0.1759 |
| Basal CORT | 4G | 2-Way ANOVA | Treatment: F (1, 31) = 7.960, p=0.0083 |
| Female:vIHKA Ctrl vs. Female:vIHKA ELS | 4G | Tukey's | q=4.300, p=0.0234, N1=10, N2=6 |
| Female:vIHKA Ctrl vs. Male:vIHKA Ctrl | 4G | Tukey's | q=3.396, p=0.0979, N1=10, N2=10 |
| Female:vIHKA Ctrl vs. Male:vIHKA ELS | 4G | Tukey's | q=4.475, p=0.0173, N1=10, N2=9 |
| Female:vIHKA ELS vs. Male:vIHKA Ctrl | 4G | Tukey's | q=1.359, p=0.7722, N1=6, N2=10 |
| Female:vIHKA ELS vs. Male:vIHKA ELS | 4G | Tukey's | q=0.312, p=0.9961, N1=6, N2=9 |
| Male:vIHKA Ctrl vs. Male:vIHKA ELS | 4G | Tukey's | q=1.170, p=0.8412, N1=10, N2=9 |
| Basal CORT vs. % Time Seizing: vIHKA CORT ∩ vIHKA EEG | 4H | Simple linear regression | Y = 1066*X + 3.656 |
| Basal CORT vs. % Time Seizing: vIHKA CORT ∩ vIHKA EEG | 4H | Pearson r correlation | r=0.8267, $R^2$=0.6834, p=0.0218 |

**Figure 5- Survival**

| Description | Figure | Statistical test | Results |
|---|---|---|---|
| Probability of survival: vIHKA Ctrl M vIHKA ELS M | 5A | Log-rank(Mantel-Cox) | χ2=0.001873, p=0.9655 |
| % change in body weight: vIHKA Ctrl M vs. vIHKA ELS M | 5B | Unpaired t test | t=0.3044, df=16, p=0.7647 |
| Probability of survival: vIHKA Ctrl F vs. vIHKA ELS F | 5C | Log-rank (Mantel-Cox) | χ2=4.102, p=0.0428 |
| % change in body weight: vIHKA Ctrl F vs. vIHKA ELS F | 5D | Unpaired t test | t=0.9035, df=15, 0.3805 |

**Figure 6- Histology**

| Description | Figure | Statistical test | Results |
|---|---|---|---|
| PVN c-Fos(+) cells (%) | 6B | 2-Way ANOVA | Interaction: F (1, 66) = 1.38 p=0.2430 |
| PVN c-Fos(+) cells (%) | 6B | 2-Way ANOVA | Sex: F (1, 66) = 15.57, p=0.0002 |
| PVN c-Fos(+) cells (%) | 6B | 2-Way ANOVA | Treatment: F (1, 66) = 2.84 p=0.0963 |
| Female:vIHKA Ctrl vs. Female:vIHK ELS | 6B | Tukey's | q=0.5281, p=0.9821, N1=1 N2=19 |
| Female:vIHKA Ctrl vs. Male:vIHKA Ctrl | 6B | Tukey's | q=3.018, p=0.1532, N1=16, N2=24 |
| Female:vIHKA Ctrl vs. Male:vIHKA E | 6B | Tukey's | q=5.061, p=0.0036, N1=16, N2=11 |
| Female:vIHKA ELS vs. Male:vIHKA | 6B | Tukey's | q=2.588, p=0.2687, N1=19, N2=24 |
| Female:vIHKA ELS vs. Male:vIHKA ELS | 6B | Tukey's | q=4.759, p=0.0068, N1=19, N2=11 |
| Male:vIHKA Ctrl vs. Male:vIHKA EL | 6B | Tukey's | q=2.769, p=0.2144, N1=24, N2=11 |

**Figure 7- Female Behavior #1**

| Description | Figure | Statistical test | Results |
|---|---|---|---|
| Homecage locomotor activity: vIHKA Ctrl F vs. vIHKA ELS F | 7A | Mixed-effects analysis | Interaction: F (5, 43) = 1.022, p=0.4166 |
| Homecage locomotor activity: vIHKA Ctrl F vs. vIHKA ELS F | 7A | Mixed-effects analysis | Time: F (2.221, 19.10) = 2.026, p=0.1559 |
| Homecage locomotor activity: vIHKA Ctrl F vs. vIHKA ELS F | 7A | Mixed-effects analysis | Treatment: F (1, 9) = 0.357 p=0.5647 |
| day0 PM | 7A | Šídák's | t=1.354, p=0.7762, N1=5, N2=6 |
| day1 AM | 7A | Šídák's | t=0.5182, p=0.9968, N1=5, N2=6 |
| day1 PM | 7A | Šídák's | t=1.027, p=0.9106, N1=5, N2=6 |
| day2 AM | 7A | Šídák's | t=0.3417, p=0.9997, N1=4, N2=6 |
| day2 PM | 7A | Šídák's | t=1.066, p=0.9050, N1=5, N2=6 |
| day3 AM | 7A | Šídák's | t=0.1229, p>0.9999, N1=5, N2=5 |
| Max startle response at 95dB: vIHKA Ctrl F vs. vIHKA ELS F | 7B | Welch's t test | t=2.208, df=11.17, p=0.049 |
| OFT- time in center: vIHKA Ctrl F vs. vIHKA ELS F | 7C | Unpaired t test | t=0.2291, df=28, p=0.8205 |
| OFT- distance in center: vIHKA Ctrl F vIHKA ELS F | 7D | Unpaired t test | t=0.3379, df=30, p=0.7378 |

| | | | |
|---|---|---|---|
| OFT - total distance traveled: vIHKA Ctrl F vs. vIHKA ELS F | 7E | Unpaired t test | t=0.3808, df=30, p=0.7060 |
| LD - time in dark zone: vIHKA Ctrl F vs. vIHKA ELS F | 7F | Unpaired t test | t=0.4718, df=30, p=0.6405 |
| LD - distance in dark zone: vIHKA Ctrl F vs. vIHKA ELS F | 7G | Unpaired t test | t=3.158, df=30, p=0.0036 |
| LD - total distance traveled: vIHKA Ctrl F vs. vIHKA ELS F | 7H | Unpaired t test | t=1.553, df=30, p=0.1309 |
| EPM - time in closed arm: vIHKA Ctrl F vs. vIHKA ELS F | 7I | Unpaired t test | t=0.3910, df=29, p=0.6986 |
| EPM - distance in closed arm: vIHKA Ctrl F vs. vIHKA ELS F | 7J | Unpaired t test | t=0.4885, df=30, p=0.6287 |
| EPM - total distance traveled: vIHKA Ctrl F vs. vIHKA ELS F | 7K | Unpaired t test | t=0.6694, df=30, p=0.5084 |

**Figure 8 - Female Behavior #2**

| Description | Figure | Statistical test | Results |
|---|---|---|---|
| SPT - avg sucrose preference: vIHKA Ctrl F vs. vIHKA ELS F | 8A | Unpaired t test | t=0.9039, df=28, p=0.3738 |
| TST - time spent immobile: vIHKA Ctrl vs. vIHKA ELS F | 8B | Unpaired t test | t=2.300, df=28, p=0.0291 |
| TST - latency to immobility: vIHKA Ctrl vs. vIHKA ELS F | 8C | Unpaired t test | t=3.370, df=23, p=0.0026 |

**Figure 9 - Male Behavior #1**

| Description | Figure | Statistical test | Results |
|---|---|---|---|
| Homecage locomotor activity: vIHKA C M vs. vIHKA ELS M | 9A | Mixed-effects analysis | Interaction: F (5, 37) = 0.2169, p=0.9531 |
| Homecage locomotor activity: vIHKA C M vs. vIHKA ELS M | 9A | Mixed-effects analysis | Time: F (5, 37) = 2.886, p=0.0268 |
| Homecage locomotor activity: vIHKA C M vs. vIHKA ELS M | 9A | Mixed-effects analysis | Treatment: F (1, 8) = 0.130 p=0.7276 |
| day0 PM | 9A | Šídák's | t=0.9029, p=0.9383, N1=7, N2=3 |
| day1 AM | 9A | Šídák's | t=0.2420, p>0.9999, N1=6, N2=3 |
| day1 PM | 9A | Šídák's | t=0.1866, p>0.9999, N1=6, N2=3 |
| day2 AM | 9A | Šídák's | t=0.3225, p=0.9997, N1=7, N2=3 |
| day2 PM | 9A | Šídák's | t=0.05353, p>0.9999, N1=6, N2=3 |
| day3 AM | 9A | Šídák's | t=0.2479, p>0.9999, N1=7, N2=3 |

| | | | |
|---|---|---|---|
| Max startle response at 95dB: vIHKA C M vs. vIHKA ELS M | 9B | Welch's t test | t=1.385, df=20.74, p=0.1809 |
| OFT - time in center: vIHKA Ctrl M vs. vIHKA ELS M | 9C | Unpaired t test | t=0.2163, df=28, p=0.8303 |
| OFT - distance in center: vIHKA Ctrl M vs. vIHKA ELS M | 9D | Unpaired t test | t=2.214, df=28, p=0.0352 |
| OFT - total distance traveled: vIHKA Ctrl M vs. vIHKA ELS M | 9E | Unpaired t test | t=0.8667, df=31, p=0.3928 |
| LD - time in dark zone: vIHKA Ctrl M vs. vIHKA ELS M | 9F | Unpaired t test | t=0.7442, df=29, p=0.4627 |
| LD - distance in dark zone: vIHKA Ctrl M vs. vIHKA ELS M | 9G | Unpaired t test | t=0.4889, df=28, p=0.6287 |
| LD - total distance traveled: vIHKA Ctrl M vs. vIHKA ELS M | 9H | Unpaired t test | t=0.9253, df=30, p=0.3622 |
| EPM - time in closed arm: vIHKA Ctrl M vs. vIHKA ELS M | 9I | Unpaired t test | t=1.118, df=25, p=0.2743 |
| EPM - distance in closed arm: vIHKA Ctrl M vs. vIHKA ELS M | 9J | Unpaired t test | t=2.068, df=28, p=0.0480 |
| EPM - total distance traveled: vIHKA Ctrl M vs. vIHKA ELS M | 9K | Unpaired t test | t=1.809, df=29, p=0.0808 |

**Figure 10- Male Behavior #2**

| Description | Figure | Statistical test | Results |
|---|---|---|---|
| SPT- avg sucrose preference: vIHKA Ctrl vs. vIHKA ELS F | 10A | Unpaired t test | t=0.2900, df=31, p=0.7738 |
| TST- time spent immobile: vIHKA Ctrl F vs vIHKA ELS F | 10B | Unpaired t test | t=0.5980, df=28, p=0.5547 |
| TST- latency to immobility: vIHKA Ctrl F vs vIHKA ELS F | 10C | Unpaired t test | t=0.6104, df=28, p=0.5465 |

**Figure S2 Supplemental Statistics**

| Description | Figure | Statistical test | Results |
|---|---|---|---|
| vIHKA EEG ∩ vIHKA Behavior: selected features, norm. | S2A | Multiple linear regression | F (11,10) = 3.67, $R^2$=0.8199, p=0.0248 |
| Immobility TST validation: DLC vs. handscore | S2F | Simple linear regression | Y = 0.7796*X + 24.54 |
| Immobility TST validation: DLC vs. handscore | S2F | Pearson r correlation | r=0.7143, $R^2$=0.5103, p<0.0001 |
| Latency TST validation: DLC vs. handscore | S2G | Simple linear regression | Y = 0.3826*X + 29.20 |
| Latency TST validation: DLC vs. handscore | S2G | Pearson r correlation | r=0.3196, $R^2$=0.1022, p=0.0367 |
| Seizure detection validation: Seizy vs. handscore | S2H | Simple linear regression | Y = 0.9921*X – 0.4646 |

| | | | |
|---|---|---|---|
| Seizure detection validation: Seizy vs. hand score | S2H | Pearson r correlation | r=0.9732, $R^2$=0.9470, p=0.0011 |

**Figure S3 Supplemental Behavior**

| Description | Figure | Statistical test | Results |
|---|---|---|---|
| LD- time in light zone: vIHKA Ctrl F vs. vIHKA ELS F | S3B | Unpaired t test | t=0.5128, df=30, p=0.611[illegible] |
| LD- distance in light zone: vIHKA Ctrl F [illegible] vIHKA ELS F | S3B | Unpaired t test | t=0.1857, df=30, p=0.853[illegible] |
| LD- immobility: vIHKA Ctrl F vs. vIHKA [illegible] F | S3B | Unpaired t test | t=1.834, df=30, p=0.0766 |
| LD- time in light zone: vIHKA Ctrl M vs. vIHKA ELS M | S3A | Unpaired t test | t=0.8292, df=29, p=0.413[illegible] |
| LD- distance in light zone: vIHKA Ctrl M [illegible] vIHKA ELS M | S3A | Unpaired t test | t=0.5581, df=30, p=0.580[illegible] |
| LD- immobility: vIHKA Ctrl M vs. vIHKA [illegible] M | S3A | Unpaired t test | t=1.175, df=30, p=0.2493 |
| OFT- time in periphery: vIHKA Ctrl F vs. vIHKA ELS F | S3D | Unpaired t test | t=0.2291, df=28, p=0.820[illegible] |
| OFT- distance in periphery: vIHKA Ctrl F [illegible] vIHKA ELS F | S3D | Unpaired t test | t=0.3693, df=30, p=0.714[illegible] |
| OFT- immobility: vIHKA Ctrl F vs. vIHKA ELS F | S3D | Unpaired t test | t=0.2374, df=30, p=0.814[illegible] |
| OFT- time in periphery: vIHKA Ctrl M vs. vIHKA ELS M | S3C | Unpaired t test | t=0.2173, df=28, p=0.829[illegible] |
| OFT- distance in periphery: vIHKA Ctrl [illegible] vs. vIHKA ELS M | S3C | Unpaired t test | t=0.7107, df=31, p=0.482[illegible] |
| OFT- immobility: vIHKA Ctrl M vs. vIHK[illegible] ELS M | S3C | Unpaired t test | t=0.7317, df=31, p=0.469[illegible] |
| EPM- time in open: vIHKA Ctrl F vs. vIH[illegible] ELS F | S3F | Unpaired t test | t=0.5288, df=28, p=0.601 |
| EPM- distance in open: vIHKA Ctrl F vs. vIHKA ELS F | S3F | Unpaired t test | t=0.9736, df=29, p=0.338[illegible] |
| EPM- immobility: vIHKA Ctrl F vs. vIHK[illegible] ELS F | S3F | Unpaired t test | t=0.2201, df=29, p=0.827[illegible] |
| EPM- time in open: vIHKA Ctrl M vs. vIH[illegible] ELS M | S3E | Unpaired t test | t=0.5231, df=23, p=0.605[illegible] |
| EPM- distance in open: vIHKA Ctrl M vs. vIHKA ELS M | S3E | Unpaired t test | t=1.039, df=26, p=0.3082 |
| EPM- immobility: vIHKA Ctrl M vs. vIHK[illegible] ELS M | S3E | Unpaired t test | t=1.753, df=28, p=0.0906 |
| ASR- max startle response (95/100/110dB) | S3G | Mixed-effects model (REML) | dB: F (2, 69) = 30.08, p<0.0001 |
| ASR- max startle response (95/100/110dB) | S3G | Mixed-effects model (REML) | Sex: F (1, 36) = 5.834, p=0.0209 |

| | | | |
|---|---|---|---|
| ASR- max startle response (95/100/110dB) | S3G | Mixed-effects model (REML) | Treatment: F (1, 36) = 1.478, p=0.2319 |
| ASR - max startle response (95/100/110dB) | S3G | Mixed-effects model (REML) | dB x Sex: F (2, 69) = 0.3003, p=0.7416 |
| ASR - max startle response (95/100/110dB) | S3G | Mixed-effects model (REML) | dB x Treatment: F (2, 69) = 0.6591, p=0.5206 |
| ASR - max startle response (95/100/110dB) | S3G | Mixed-effects model (REML) | Sex x Treatment: F (1, 36) = 1.640, p=0.2085 |
| ASR - max startle response (95/100/110dB) | S3G | Mixed-effects model (REML) | Interaction: F (2, 69) = 1.271, p=0.2870 |
| ASR - avg startle response (95/100/110dB) | S3H | Mixed-effects model (REML) | dB: F (2, 68) = 32.92, p<0.0001 |
| ASR - avg startle response (95/100/110dB) | S3H | Mixed-effects model (REML) | Sex: F (1, 36) = 7.108, p=0.0114 |
| ASR - avg startle response (95/100/110dB) | S3H | Mixed-effects model (REML) | Treatment: F (1, 36) = 2.279, p=0.1398 |
| ASR - avg startle response (95/100/110dB) | S3H | Mixed-effects model (REML) | dB x Sex: F (2, 68) = 0.5863, p=0.5592 |
| ASR - avg startle response (95/100/110dB) | S3H | Mixed-effects model (REML) | dB x Treatment: F (2, 68) = 0.8885, p=0.4160 |
| ASR - avg startle response (95/100/110dB) | S3H | Mixed-effects model (REML) | Sex x Treatment: F (1, 36) = 1.608, p=0.2128 |
| ASR - avg startle response (95/100/110dB) | S3H | Mixed-effects model (REML) | Interaction: F (2, 68) = 0.6904, p=0.5048 |

**Figure S4 2way Acute EEG**

| **Description** | **Figure** | **Statistical test** | **Results** |
|---|---|---|---|
| Epileptiform activity (%) | S4A | 2-Way ANOVA | Interaction: F (1, 24) = 36.37, p<0.0001 |
| Epileptiform activity (%) | S4A | 2-Way ANOVA | Sex: F (1, 24) = 4.996, p=0.0350 |
| Epileptiform activity (%) | S4A | 2-Way ANOVA | Treatment: F (1, 24) = 4.946, p=0.0358 |
| Female:Control vs. Female:ELS | S4A | Tukey's | q=8.212, p<0.0001, N1=6, N2=8 |
| Female:Control vs. Male:Contro | S4A | Tukey's | q=3.666, p=0.0709, N1=6, N2=7 |
| Female:Control vs.Male:ELS | S4A | Tukey's | q=0.01070, p>0.9999, N1=6, N2=7 |
| Female:ELS vs. Male:Control | S4A | Tukey's | q=4.629, p=0.0158, N1=8, N2=7 |
| Female:ELS vs. Male:ELS | S4A | Tukey's | q=8.581, p<0.0001, N1=8, N2=7 |
| Male:Control vs. Male:ELS | S4A | Tukey's | q=3.826, p=0.0559, N1=7, N2=7 |
| % Time seizing | S4B | 2-Way ANOVA | Interaction: F (1, 29) = 5.508, p=0.0260 |
| % Time seizing | S4B | 2-Way ANOVA | Sex: F (1, 29) = 0.07101, p=0.7918 |
| %Time seizing | S4B | 2-Way ANOVA | Treatment: F (1, 29) = 0.005922, p=0.9392 |

| | | | |
|---|---|---|---|
| Female:Control vs. Female:ELS | S4B | Tukey's | q=2.303, p=0.3790, N1=9, N2=8 |
| Female:Control vs. Male:Control | S4B | Tukey's | q=2.111, p=0.4548, N1=9, N2=8 |
| Female:Control vs. Male:ELS | S4B | Tukey's | q=0.3484, p=0.9946, N1=9, N2=8 |
| Female:ELS vs. Male:Control | S4B | Tukey's | q=0.1869, p=0.9992, N1=8, N2=8 |
| Female:ELS vs. Male:ELS | S4B | Tukey's | q=2.577, p=0.2838, N1=8, N2=8 |
| Male:Control vs. Male:ELS | S4B | Tukey's | q=2.390, p=0.3470, N1=8, N2=8 |
